# DYRK1A regulates cancer cell and cancer-associated fibroblast secretomes to foster an immunosuppressive microenvironment in pancreatic cancer

**DOI:** 10.1101/2025.11.26.690756

**Authors:** Silvia Pascual-Sabater, Sushmita Varhadi, Chiara Di Vona, Marta Celma, Cristina Fillat, Susana de la Luna

## Abstract

Pancreatic ductal adenocarcinoma (PDAC) is characterized by a desmoplastic, immunosuppressive stroma dominated by cancer-associated fibroblasts (CAFs). Paracrine signals from cancer cells and CAFs shape the tumor microenvironment (TME), facilitating interactions that drive tumor growth and immune evasion, yet the molecular regulators of this crosstalk remain incompletely understood. Here, we show that Dual-specificity Tyrosine-Regulated Kinase 1A (DYRK1A), previously shown to promote PDAC growth by stabilizing c-MET in cancer cells, is also expressed in CAFs at levels comparable to those in cancer cells. Secretome profiling of DYRK1A-depleted cancer cells and CAFs by quantitative mass spectrometry revealed DYRK1A-dependent factors in each cell type associated with terms related to cell migration, while combining the DYRK1A-regulated proteins from both compartments highlighted additional categories associated with immune infiltration and cell-cell interactions in the TME. Genetic or pharmacological inhibition of DYRK1A reduced CCL2, CCL5 and CSF-1 secretion in cancer cells, and CXCL12 in CAFs, linking DYRK1A activity to immunosuppressive paracrine signaling. Conditioned media from DYRK1A-depleted cancer cells or CAFs impaired the migration of monocytes and myeloid-derived suppressor cells, indicating that DYRK1A remodels the PDAC secretome to recruit immunosuppressive myeloid populations. These findings extend the established role of DYRK1A beyond cancer cell-intrinsic signaling, highlighting its role as a regulator of secreted factors that shape the PDAC TME, and position this kinase as a potential target to reduce immunosuppressive signaling.

**Significance:** DYRK1A regulates chemokine and cytokine secretion in PDAC cancer cells and CAFs, promoting recruitment of immunosuppressive myeloid populations, and revealing a previously unrecognized role for this kinase in shaping the TME.

## Introduction

Pancreatic ductal adenocarcinoma (PDAC) is a highly aggressive malignancy with an incidence rising by about 1% per year and a 5-year survival rate of only 13% (1). Although advances in chemotherapy regimens have improved outcomes in patients with resectable disease (2), more than 80% of patients are diagnosed at advanced stages, precluding surgical resection (3). Moreover, PDAC tumors are notoriously resistant to chemotherapy, targeted therapies, and immunotherapy, underscoring the urgent need for novel therapeutic strategies (4).

PDAC is characterized by a complex tumor microenvironment (TME) with a dense stromal matrix composed of diverse cellular and acellular elements (4). Cancer-associated fibroblasts (CAFs) are the most abundant stromal cell type and actively contribute to tumor progression by secreting growth factors, inflammatory mediators, and extracellular matrix (ECM) proteins that promote cancer cell proliferation, invasion, therapy resistance, and immune evasion (5). Together with cancer cells, CAFs shape an immunosuppressive TME dominated by myeloid cell infiltration and typically devoid of CD4^+^ or CD8^+^ T cells, rendering PDAC an immunologically ’cold’ tumor (4). Anti-stromal therapies for PDAC have shown limited efficacy, likely reflecting the complex biology of tumor-stroma interactions (4). Notably, cancer cells and CAFs share signaling pathways that regulate immune cell recruitment and function, highlighting the potential of targeting shared molecular regulators to modulate TME composition.

Dual-specificity tyrosine-regulated kinases (DYRKs) are an evolutionarily conserved family of CMGC group kinases within the eukaryote kinome (6). In humans, the class I paralogues DYRK1A and DYRK1B have been implicated in several malignancies due to their broad cellular functions (7). Both kinases are overexpressed in PDAC, where they promote tumor cell proliferation and survival (8–10). We have previously demonstrated that DYRK1A is upregulated in PDAC from preneoplastic stages and promotes tumor growth by stabilizing the receptor tyrosine kinase (RTK) c-MET in cancer cells (9). However, whether DYRK1A also regulates paracrine communication within the PDAC TME remains unknown.

Here, we show that, unlike its paralogue DYRK1B, DYRK1A is expressed at comparable levels in PDAC cancer cells and CAFs, where it is overexpressed compared with normal fibroblasts. Using patient-derived PDAC CAFs and cancer cell lines with reduced DYRK1A expression and/or activity, we characterized their DYRK1A-dependent secretome by quantitative label-free mass spectrometry. We identified distinct yet functionally convergent alterations in secreted proteins from both cell types that modulate immune infiltrates. Specifically, DYRK1A depletion in cancer cells diminished secretion of CCL2/MCP-1, CCL5/RANTES, and CSF-1/M-CSF, resulting in reduced migration of both monocytes and myeloid-derived suppressor cells (MDSCs). Moreover, DYRK1A silencing or inhibition in CAFs reduced CXCL12/SDF-1 secretion and substantially impaired the recruitment of MDSCs, with a milder effect observed on monocyte migration. Together, these findings reveal a previously unrecognized role for DYRK1A in regulating stromal–cancer cell crosstalk through secreted mediators and highlight its potential as a therapeutic target to reprogram the PDAC TME.

## Materials and Methods

### Human specimens

PDAC samples were obtained from treatment-naïve patients undergoing surgery at Hospital Clínic de Barcelona. All patients provided written informed consent prior to sample collection. The study was approved by the Hospital Clínic Ethical Committee and conducted in accordance with the Declaration of Helsinki.

### Chemicals and reagents

The sources of chemicals and reagents (supplier and reference) are listed in Supplementary Table S1, plasmids in Table S2, oligonucleotides in Table S3 and antibodies in Table S4.

### Cell culture

PDAC cell lines PANC-1 (RRID:CVCL_0480), MIA PaCa-2 (RRID:CVCL_0428), Hs 766T (RRID:CVCL_0334), and derived cell lines were grown in Dulbecco’s Modified Eagle’s medium (DMEM) supplemented with 10% fetal bovine serum (FBS) and 100 units/ml penicillin and 100 µg/ml streptomycin. For HEK293FT (RRID:CVCL_6911) and immortalized PDAC CAFs, FBS was heat inactivated and media further supplemented with 2 mM L-glutamine. All cell lines were maintained at 37°C in a humidified atmosphere with 5% CO_2_ and routinely tested for mycoplasma contamination. Cell authentication was performed by short tandem repeat profiling. DYRK inhibitors harmine and cirtuvivint (SM08502) were dissolved in dimethyl sulfoxide (DMSO). Control cells received the corresponding DMSO concentration.

### CRISPR-Cas9-mediated DYRK1A and DYRK1B knockout (KO)

PANC-1 and MIA PaCa-2 cells were transfected with CRISPR-Cas9 constructs targeting DYRK1A or DYRK1B. Two sgRNAs were designed for each kinase (DYRK1A-sgRNA1: 5’-CAGGCAGCTGGGCATATTA-3’; DYRK1A-sgRNA2: 5’-TCCATGTAGCATACCATGTA-3’; DYRK1B-sgRNA1: 5’-TATGTGAATAGTGCCAGGCG-3’; DYRK1B-sgRNA2: 5’-GGTTACTGAGTGCTCACCAA-3’) and cloned into pX458_pSpCas9(BB)-2A-GFP. Transfections were performed with Lipofectamine 3000, and 48 h later, GFP-positive cells were sorted by flow cytometry (Becton Dickinson, RRID:SCR_019593) and seeded at one cell/well in 96-well plates. Medium was replaced every 6-7 days until clones reached confluency. Clones were screened by PCR genotyping using primer pairs that detected deleted and wild-type alleles (Supplementary Table S3) and validated by Western blotting.

### Isolation and culture of primary pancreatic fibroblasts

CAFs and normal fibroblasts (NFs) were isolated from human PDAC tissues or adjacent pancreatic tissues, respectively, as described (11) with minor modifications. Details are provided in Supplementary Materials and Methods. Primary PDAC CAFs were immortalized with a lentiviral vector encoding the SV40 large T-antigen (pLOX-Ttag-iresTK). Immortalized CAFs were selected based on outgrowth after passaging and maintained as polyclonal populations. For DYRK1A knockdown (KD), immortalized CAFs were transduced with pLKO.1-puro-based lentiviral vectors (Sigma Mission shRNA collection), including a non-targeting control (SHC002, shCTRL) and two DYRK1A-targeting shRNAs (shD1A#1, TRCN0000022999; shD1A#2, TRCN0000199188). Cells were selected 48 h post-transduction with 3 μg/mL puromycin for 3-4 days, and KD efficiency was confirmed by RT-qPCR and/or Western blotting.

### Human monocyte isolation and monocytic MDSC generation

Peripheral blood monocytes were isolated from buffy coats of healthy donors (Barcelona Public Blood and Tissue Bank) using a Ficoll gradient (Ficoll-Paque Plus density 1.077 g/mL) followed by negative selection. See Supplementary Materials and Methods for details. Monocytes were differentiated into MDSC by culturing with 10 ng/mL IL-6 and 10 ng/mL GM-CSF for 6 days (12).

### Generation of conditioned media (CM)

Cells at 80% confluency were washed three times with PBS and incubated with serum-free DMEM. CM was collected after 24 h (cancer cells) or 48 h (CAFs), centrifuged (300x*g*, 10 min, 4°C), and stored in single-use aliquots at −70°C. For mass spectrometry, CM was concentrated using Amicon^TM^ Ultra-15 10-kDa filters.

### Migration assays

Cell migration was assessed using polycarbonate Transwell inserts (5-µm pores; 24-well plates; Falcon® #353097). Cells (5 x 10^5^) in serum-free medium were seeded in the upper chamber, and 800 μL of CM was added to the lower chamber. Cells were allowed to migrate for 5 h (monocytes) or 15 h (MDSCs) at 37°C. For neutralization experiments, CM was pre-incubated for 30 min at 37°C with antibodies to CCL2 or CXCL12 (Supplementary Table S4) before being added to the lower chamber. Migrated cells were collected from the lower chamber and quantified by flow cytometry using CountBright^TM^ Absolute Counting Beads, with 10,000 beads acquired on a BD LSRFortessa SORP 4L (BD Biosciences, RRID:SCR_018655). Data were analyzed with FlowJo™ v10.

### RNA and protein expression analysis

Procedures for RNA quantification as well as protein identification and quantification by ELISA, Western blotting, and immunohistochemistry are described in the Supplementary Materials and Methods.

### Secretome analysis by liquid chromatography-mass spectrometry (LC-MS)

Identification of proteins by LC-MS in the CM was performed at the CRG/UPF Proteomics Unit. Details on sample preparation and chromatographic and MS analysis are included in Supplementary Materials and Methods. Label-free quantification was performed with four replicates per condition. Proteins with > 3 peptide-spectrum matches were retained. Protein amounts were estimated by averaging the three most intense peptides and normalized to the total protein amount in each sample. Abundances were log_2_-transformed and fold changes (FC) and *p*-values (unpaired two-tailed Student’s *t*-test) calculated. Proteins were considered deregulated if *p* ≤ 0.05 and log_2_(FC) ≤ −0.5 (downregulated in DYRK1A^KO^/KD cells) or log_2_(FC) ≥ 0.5 (upregulated in DYRK1A^KO^/KD cells) in comparison with their respective controls.

Proteins detected in ≥ 3 control replicates but ≤ 1 replicates of DYRK1A^KO^/KD cells were considered as downregulated; the inverse was used to define upregulated proteins. In these cases, *p*-value and log_2_(FC) were calculated with missing values imputed using the lowest normalized detected intensity.

### Computational tools and statistical analysis

A full list of computational tools is provided in Supplementary Materials and Methods and Table S5. All statistical analyses were performed with Prism v8.0.1. Normality was assessed using the Shapiro-Wilk test. Statistical significance in pairwise comparisons was determined with two-tailed unpaired Student’s *t*-tests, or one sample t-tests in experiments normalized to control values of 1. Differences were considered significant at *p* ≤ 0.05. Data are plotted as mean ± SEM with individual values displayed.

### Data availability

Proteomics data generated in this study are publicly available at the ProteomeXchange Consortium via the PRIDE partner repository under dataset identifier PXD022966. Summary data are available within the Article and Supplementary data files. Expression datasets from Gene Expression Omnibus (GEO) analyzed are listed in Supplementary Table S6.

## Results

### DYRK1A is overexpressed in PDAC CAFs

Given the crucial contribution of the stroma to pancreatic tumors, we hypothesized that DYRK1A might also play a role in CAFs. To explore this, we first analyzed publicly available RNA-seq data from microdissected PDAC tissues (13, 14). *DYRK1A* expression was comparable between the epithelial and stromal compartments (Fig. 1A and B). We next examined DYRK1A expression by immunohistochemistry in primary PDAC tumors. Consistent with our previous study (9), strong DYRK1A staining was observed in the epithelial compartment; notably, fibroblasts within the surrounding stroma also exhibited prominent DYRK1A expression (Fig. 1C). Moreover, CAFs isolated from PDAC tumors displayed DYRK1A protein levels comparable to those of PANC-1 cancer cells and higher than those observed in patient-matched fibroblasts from adjacent normal pancreatic tissue (Fig. 1D). In contrast, the paralogous kinase DYRK1B was readily detected in PANC-1 cells but was barely detectable in CAFs, where *DYRK1B* transcript levels were markedly lower than those of DYRK1A (Fig. 1E; Supplementary Fig. S1A). This pattern was confirmed in independent publicly available transcriptomic datasets, where *DYRK1B* expression was consistently lower than *DYRK1A* in CAFs and reduced in stromal relative to epithelial compartments (Supplementary Fig. S1B-S1D). A similar trend was observed in fibroblasts from other tumor types, including liver, colon, and breast (Supplementary Fig. S1E). Together, these results suggest that DYRK1A –but not DYRK1B– may contribute to the protumorigenic features of PDAC CAFs.

**Figure 1.**
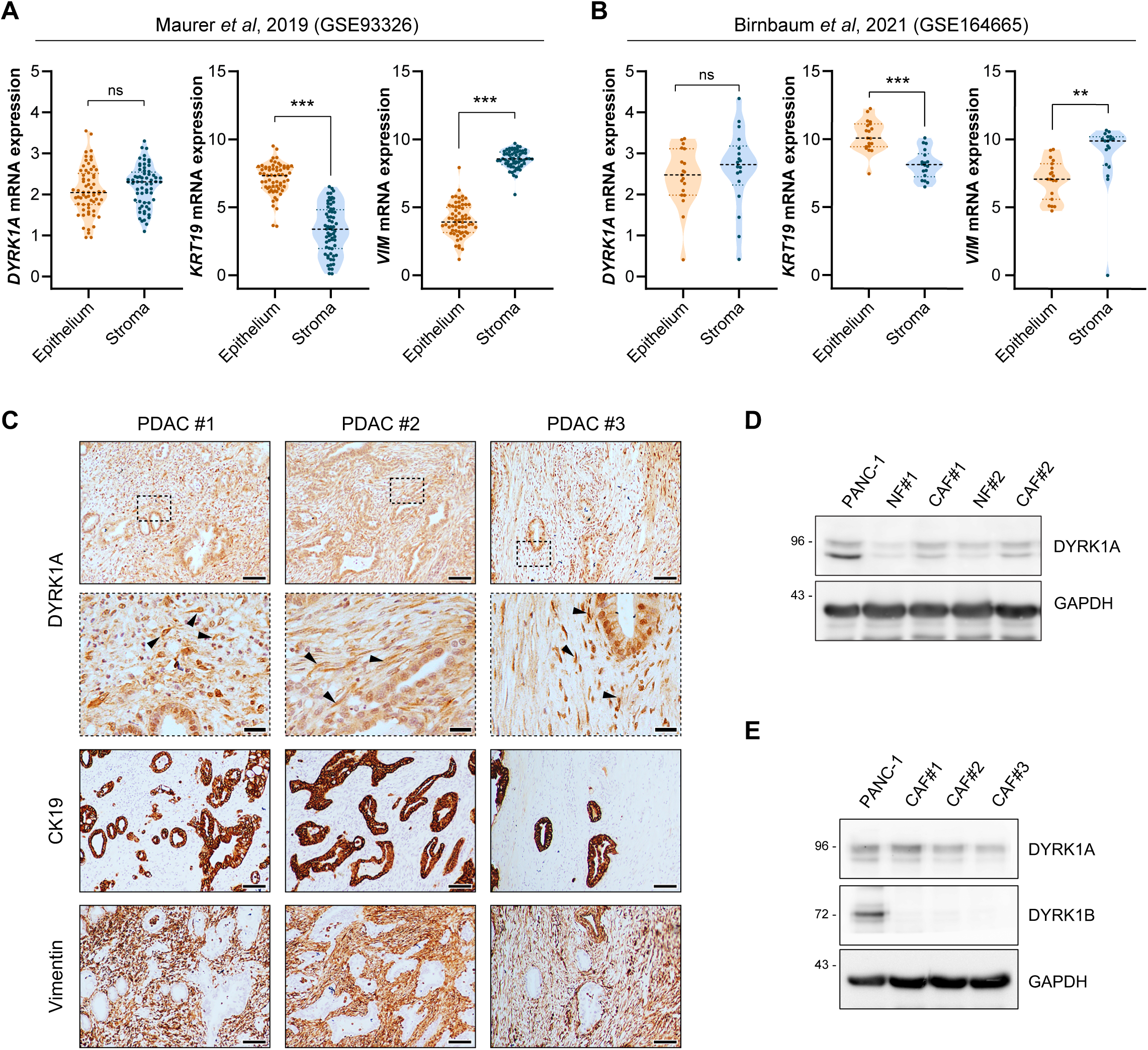
DYRK1A is expressed in both cancer cells and CAFs in PDAC tumors. **A** and **B**, Violin plots showing *DYRK1A* mRNA expression (log_2_ [TPM+1]) in matched epithelial and stromal compartments of microdissected PDAC tumors (GSE93326, *n* = 64; GSE164665, *n* = 18). *KRT19* and *VIM* were used as markers for compartment identification (*ns* = not significant, \*\**p* ≤ 0.01, \*\*\**p* ≤ 0.001; paired t-test). **C,** Immunohistochemical staining of DYRK1A on primary PDAC tumors. Cytokeratin 19 (CK19) staining highlights epithelial ductal cells, while vimentin staining identifies stromal areas. Arrowheads in higher magnification images point to representative DYRK1A^+^ CAFs. Scale bar: 100 μm (low magnification) or 25 μm (high magnification). **D,** Western blot analysis of DYRK1A expression in PANC-1 cells and primary human normal fibroblasts (NFs) or CAFs. **E,** DYRK1A and DYRK1B expression in PANC-1 cells and CAFs derived from 3 different PDAC tumors. GAPDH was used as a loading control in **D** and **E**.

### DYRK1A regulates the secretome of PDAC cancer cells

Proteins secreted by cancer cells and CAFs critically shape the PDAC TME (15). Given DYRK1A the expression in both cell types, we investigated whether DYRK1A influences the PDAC TME by modulating their secretome. DYRK1A-deficient PANC-1 cells (D1A^KO^) were generated using CRISPR/Cas9 (Supplementary Fig. S2A), and loss of DYRK1A expression was confirmed by western blotting (Supplementary Fig. S2B). D1A^KO^ clones exhibited reduced cell proliferation compared with wild-type (WT) cells (Supplementary Fig. S2C), consistent with our previous transient DYRK1A silencing results (9). Label-free MS analysis was performed on CM collected from two PANC-1 D1A^KO^ clones and WT cells 24 h after serum starvation (Fig. 2A). The number of secreted proteins detected was similar between WT and D1A^KO^ cells (Supplementary Fig. S2D), indicating that DYRK1A loss does not broadly affect overall protein secretion. Both upregulated and downregulated proteins were identified in D1A^KO^ CM (Supplementary Fig. S2E; Table S7), with substantial overlap between clones (174 shared proteins), particularly among downregulated proteins (Fig. 2B).

**Figure 2.**
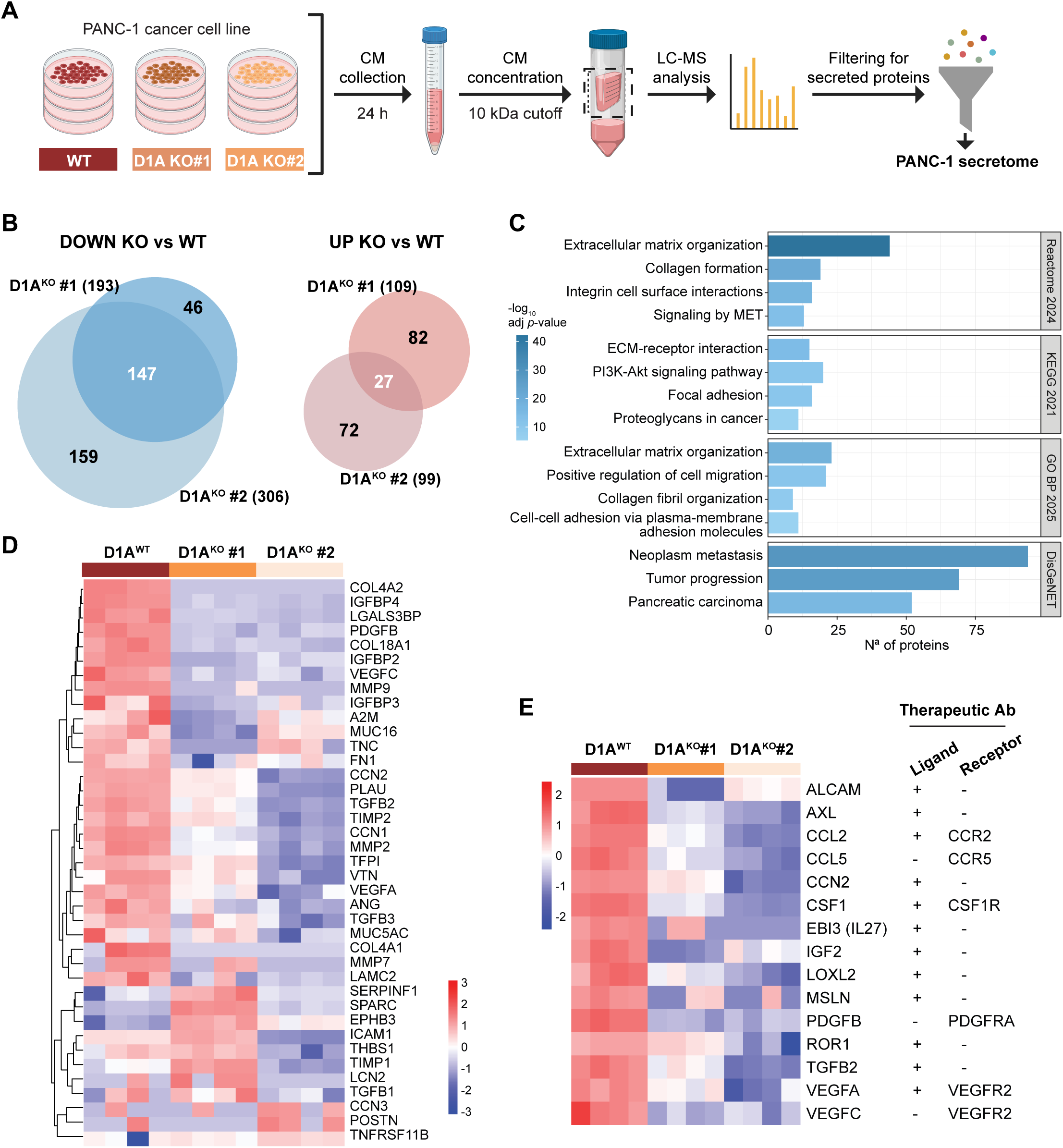
The DYRK1A-dependent secretome of PDAC cancer cells. A, Schematic representation of the MS workflow used to characterize the secretome of PANC-1 WT cells and the D1A^KO^-derived clones (D1A^KO^#1, D1A^KO^#2). LC-MS, liquid chromatography-mass spectrometry. **B,** Venn diagrams showing secreted proteins significantly down- or upregulated in PANC-1 D1A^KO^ clones compared with WT cells (*p* ≤ 0.05, −0.5 ≥ log_2_(FC) ≥ 0.5). Extended data are available in Table S7. **C,** Functional enrichment analysis of secreted proteins commonly downregulated in both PANC-1 D1A^KO^ clones (147 proteins). Selected terms are grouped into categories according to Enrichr (Reactome, Reactome Pathways 2024; KEGG 2021, Kyoto Encyclopedia of Genes and Genomes 2021; GO BP 2025, Gene Ontology Biological Process 2025 and DisGeNET). Extended data are available in Table S8. **D** and **E,** Heatmaps showing differentially secreted proteins between PANC-1 WT cells and D1A^KO^ clones, either identified as PDAC biomarkers (**D**) or associated with existing therapeutic antibodies and targeting either the ligand or the receptor as indicated (**E**). An extended overview of these proteins is provided in Supplementary Fig. S2H. Extended data for PDAC biomarkers are available in Table S9.

Functional enrichment of downregulated proteins in PANC-1 D1A^KO^ cells highlighted pathways associated with “pancreatic carcinoma”, “extracellular matrix organization”, and “positive regulation of cell migration” (Fig. 2C; Table S8), with a core subset common to these terms (Supplementary Fig. S2F). These proteins included matrix-remodeling and chemotactic factors, indicative of a DYRK1A-dependent cancer cell secretome that supports invasion and stromal recruitment. In contrast, only a few categories reached significance among upregulated proteins (Supplementary Fig. S2G). Several downregulated ECM-associated proteins –AGRN, LAMC2, CD109 and MMP9– are typically upregulated during PDAC progression and correlate with poor prognosis (16), suggesting that DYRK1A promotes the secretion of pro-tumorigenic ECM components. This effect may be reinforced by the concomitant reduction in ligand-receptor pairs, such as AGRN-LRP4 and LAMC2-ITGB1 (17, 18), detected in the DYRK1A-depleted PANC-1 secretome (Table S7). Notably, multiple downregulated proteins are reported as PDAC biomarkers (19, 20) (Fig. 2D; Table S9). Furthermore, interrogation of the therapeutic antibody database Thera-SAbDab (21) identified downregulated secreted proteins such as PDGFB, VEGFA, CCL2, CCL5, and CSF-1 as direct or indirect targets of approved or investigational antibodies (Fig. 2E; Supplementary Fig. S2H), underscoring the presence of clinically relevant factors within the PANC-1 DYRK1A-dependent secretome. Finally, several RTKs were detected in the PANC-1 CM (Supplementary Fig. S2I; Table S10), likely reflecting ectodomain shedding. As DYRK1A stabilizes VEGFR2, EGFR, and c-MET (9, 22, 23), the reduced abundance of additional RTKs in D1A^KO^ CM (Supplementary Fig. S2I), suggests a broad effect on RTK expression, stability and/or shedding. Together, these findings indicate that DYRK1A loss disrupts cancer cell secretory programs in cell–matrix and tumor–stroma interactions.

### DYRK1A-dependent secretome of PDAC CAFs

We next examined whether DYRK1A shapes the CAF secretome. CAF lines derived from primary PDAC tumors (CAF#1-3) were transduced with lentiviral vectors expressing either a non-targeting shRNA or shRNAs specific for DYRK1A (Supplementary Fig. S3A). Effective knockdown (KD) was confirmed at the RNA and protein levels (Supplementary Fig. S3B and S3C). CM collected 48 h after starvation from shCTRL and shD1A#1 CAFs was analyzed by label-free MS (Fig. 3A). The overall detection of secreted proteins was consistent across conditions (∼470 proteins; Supplementary Fig. S3D). However, the number and identity of proteins altered by DYRK1A KD varied among CAF lines (Fig. 3B and C; Table S7), likely reflecting interpatient heterogeneity. In total, 86 proteins were significantly altered in at least two CAFs and 22 were consistently deregulated in all lines (Fig. 3C; Table S7). Functional enrichment analysis showed that upregulated factors were primarily associated with ECM remodeling (Supplementary Fig. S3E), whereas downregulated proteins were enriched in categories related to cell migration and paracrine signaling, including “T-cell modulation in pancreatic cancer” and “neovascularization processes” (Fig. 3D). Notably, CXCL12 emerged as the only factor common to these categories that was consistently downregulated in all shD1A CAF lines. CXCL12 is a well-established promoter of PDAC progression, MDSC recruitment, and T-cell exclusion (4). Moreover, TheraSabDab analysis identified the CXCL12 receptor CXCR4 as a therapeutic antibody target (Supplementary Fig. S3F), further highlighting CXCL12 as functionally relevant DYRK1A-regulated CAF-derived factor.

**Figure 3.**
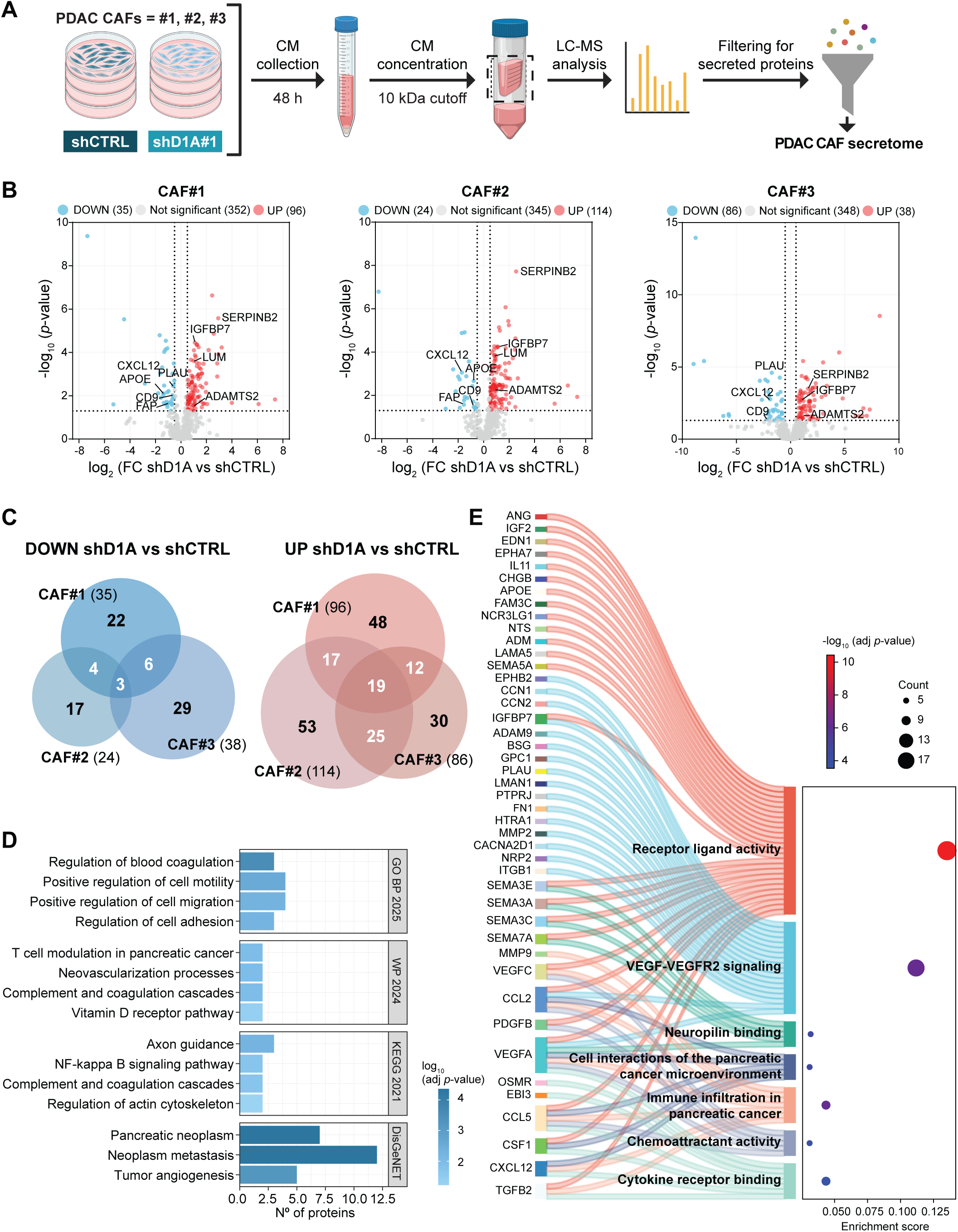
The DYRK1A-dependent secretome of PDAC CAFs. A, Schematic representation of the MS workflow used to characterize DYRK1A-dependent secretomes in CAFs derived from primary tumors of three PDAC patients (CAF#1, CAF#2, CAF#3). **B,** Volcano plots showing differentially secreted proteins between shCTRL and shDYRK1A (shD1A#1) PDAC CAFs, with significance (–log10 p-value) plotted against differences in protein abundance [log2(FC)]. The dotted line along the *y*-axis indicates significance (*p* ≤ 0.05), while the vertical dotted lines indicate fold change thresholds (log_2_(FC) ≥ 0.5 or ≤ −0.5). Extended data is available in Table S7. **C,** Venn diagrams showing secreted proteins significantly up- or downregulated in shD1A#1 CAFs compared with shCTRL (*p* ≤ 0.05, −0.5 ≥ log_2_(FC) ≥ 0.5). **D,** Functional enrichment analysis of secreted proteins significantly downregulated by DYRK1A silencing (shared by at least two CAF lines, 13 proteins). Selected terms according to Enrichr from GO BP 2025, WikiPathways Human 2024 (WP 2024), KEGG 2021 and DisGeNET are shown. Extended data are available in Table S8. **E,** Sankey plot showing the association between downregulated secreted proteins in PANC-1-derived D1A^KO^ clones and DYRK1A-depleted CAF lines, and their enriched functional categories. Links represent protein-to-pathway assignments.

Comparison of the cancer cell and CAF secretomes revealed minimal overlap: none of the upregulated proteins and only three of the downregulated ones –BGN, PLAU and SPOCK1–were commonly altered, consistent with intrinsic differences in the secretory profiles of the two cell types (15), but also suggestive of cell type-specific DYRK1A functions. Nonetheless, both DYRK1A-dependent secretomes were enriched in migration-related processes (Fig. 2C and 3D). Moreover, enrichment analysis on the combined list of downregulated proteins from both DYRK1A-depleted cell types revealed enrichment in intercellular communication categories, including “Immune infiltration in pancreatic cancer” and “Cell interactions of the pancreatic cancer microenvironment” (Fig. 3E; Supplementary Fig. S3G). Altogether, DYRK1A depletion in both PDAC cancer cells and CAFs resulted in a coordinated reduction of clinically relevant secreted factors, suggesting that DYRK1A sustains a secretory network enriched in pro-tumorigenic and druggable mediators.

### CCL2, CCL5 and CSF-1 are DYRK1A-regulated paracrine factors in PDAC cancer cells

Given their established roles in cell migration and in shaping the immunosuppressive PDAC TME (4), we focused on CCL2, CCL5 and CSF-1 in PANC-1 cells, and CXCL12 in CAFs (Fig. 3E), as relevant DYRK1A-dependent secreted factors mediating a potential immunomodulatory function of DYRK1A. Reduced secretion of CCL2 and CCL5 was confirmed by ELISA in CM from PANC-1 D1A^KO^ clones (Fig. 4A), whereas CSF-1 levels were below detection (Supplementary Fig. S4A). Consistent with the secretome results, *CCL2* and *CSF1* mRNA levels were significantly decreased in PANC-1 D1A^KO^ clones, whereas *CCL5* transcript levels remained largely unchanged (Fig. 4B), indicating that distinct regulatory mechanisms underlie DYRK1A-dependent regulation of these factors. To extend these observations to another PDAC model, we generated D1A^KO^ MIA PaCa-2 cells, which express CCL5 and CSF-1 but not CCL2 (Supplementary Fig. S4A and S4B). As observed in PANC-1, CCL5 secretion was strongly reduced in MIA PaCa-2 D1A^KO^ clones and CSF-1 secretion decreased in one clone (Fig. 4C). Treatment of three PDAC cell lines (PANC-1, MIA PaCa-2 and Hs766T) with the DYRK1 inhibitors harmine or cirtuvivint led to a general reduction in CCL2, CCL5 and CSF-1 secretion (Fig. 4D), supporting the dependence of this effect on DYRK1A kinase activity. Inhibitor-induced changes in *CCL2*, *CCL5*, and *CSF1* mRNA mirrored those observed upon DYRK1A depletion (Supplementary Fig. S4C). Among the three cell lines, PANC-1 showed a milder inhibitor response (Fig. 4D), likely reflecting co-expression of DYRK1A and DYRK1B (Supplementary Fig. S4D). Indeed, DYRK1B loss has been reported to activate *CCL5* transcription in murine PDAC cells (10), consistent with the *CCL5* mRNA increase observed upon DYRK1 inhibition in PANC-1 cells (Supplementary Fig. S4C). To assess the contribution of DYRK1B, we generated PANC-1 D1B^KO^ cells (Supplementary Fig. S4E and S4F). In contrast to DYRK1A loss, DYRK1B depletion caused significant upregulation of *CCL2* and *CCL5* (Supplementary Fig. S4G). Furthermore, analysis of the TCGA PDAC dataset revealed a positive correlation between *DYRK1A* transcript levels and those of *CCL2*, *CCL5* and *CSF1*, whereas *DYRK1B* showed no such correlation (Supplementary Fig. S4H and S4I), supporting distinct regulatory roles for the two paralogues.

**Figure 4.**
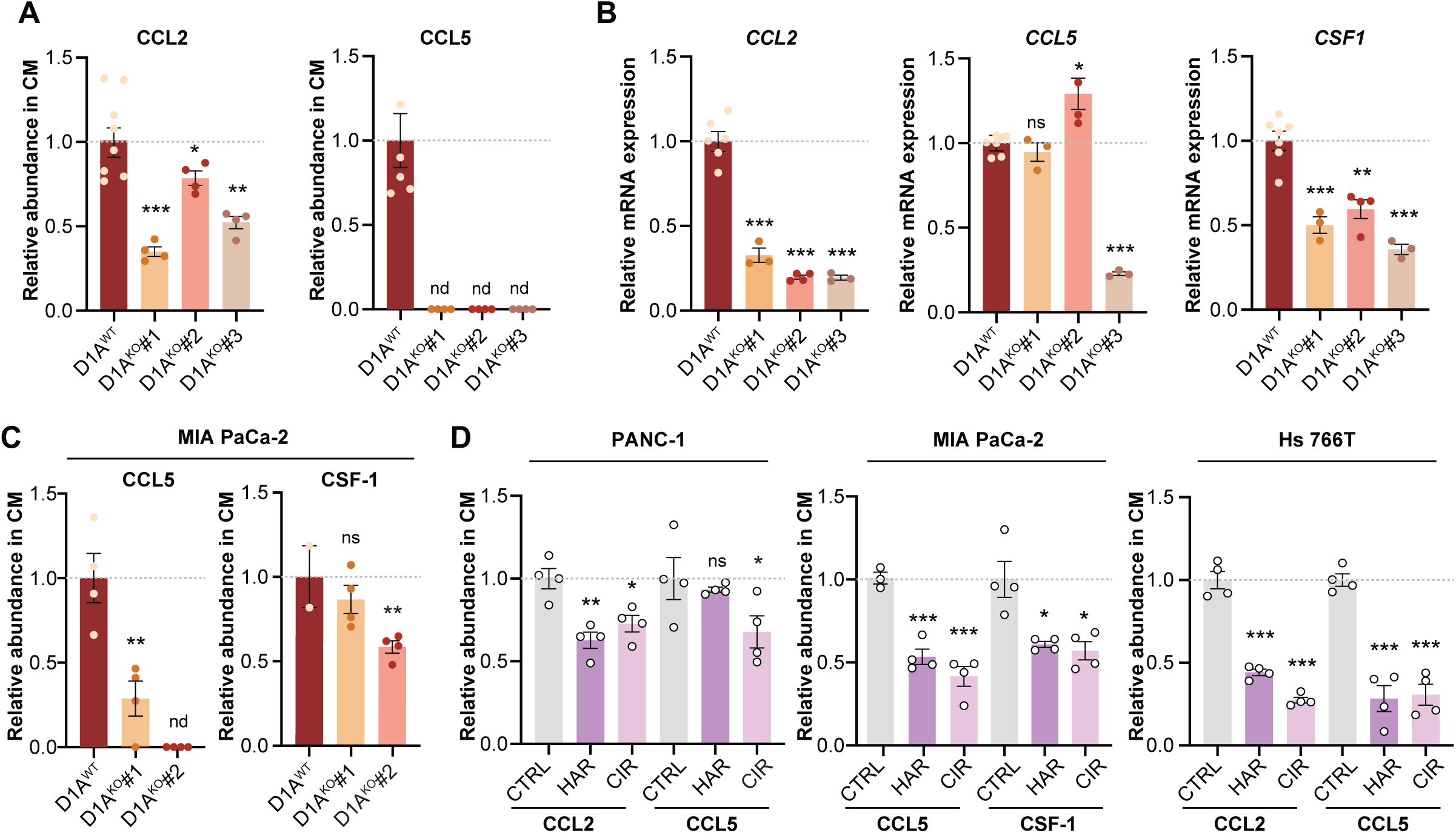
Genetic depletion or pharmacological inhibition of DYRK1A reduces secretion of the cytokines CCL2, CCL5 and CSF-1. A, ELISA-based quantification of CCL2 and CCL5 by ELISA in the CM of PANC-1 WT cells and D1A^KO^ clones cultured in serum-free conditions. Data are expressed relative to parental PANC-1 cells (WT) and represented as mean ± SEM. **B,** mRNA expression of *CCL2*, *CCL5* and *CSF-1* analyzed by RT-qPCR in PANC-1 WT cells and D1A^KO^ clones. *HPRT1* was used as a housekeeping gene. Data are expressed relative to PANC-1 WT cells values and shown as mean ± SEM. **C,** ELISA-based quantification of CCL5 and CSF-1 levels in the CM of MIA PaCa-2 WT cells and D1A^KO^ clones grown in the absence of serum. Data are expressed relative to MIA PaCa-2 WT cells and shown as mean ± SEM. **D,** Levels of the indicated cytokines in the CM of PANC-1, MIA PaCa-2 and Hs 766T treated for 24 h in serum-free conditions with DMSO as vehicle (CTRL), 10 µM harmine (HAR) or 1 µM cirtuvivint (CIR) quantified by ELISA. Values are expressed relative to vehicle controls and represented as mean ± SEM. In all panels: *nd* = not detected; *ns* = not significant, \**p* ≤ 0.05, \*\**p* ≤ 0.01, \*\*\**p* ≤ 0.001; unpaired *t*-test.

Altogether, these findings establish DYRK1A as a critical regulator of the expression and secretion of CCL2, CCL5 and CSF-1, linking its activity to immunosuppressive paracrine networks.

### CXCL12 is a DYRK1A-regulated paracrine factor in PDAC CAFs

We next examined whether DYRK1A activity regulates CXCL12 secretion in CAFs. DYRK1A depletion reduced CXCL12 secretion in CAFs (Fig. 5A; Supplementary Fig. S5A), accompanied by decreased mRNA expression (Fig. S5B), pointing to a transcriptional mechanism. This effect was recapitulated in CAFs from all three patients treated with harmine or cirtuvivint, with the latter inducing a stronger reduction (Fig. 5C; Supplementary Fig. S5B), indicating that CXCL12 expression depends on DYRK1A kinase activity. DYRK1B is minimally expressed in CAFs (Fig. 1E), ruling out confounding paralogue effects. Analysis of TCGA PDAC data and a stroma-specific dataset (GSE93326) revealed a positive correlation between *CXCL12* and *DYRK1A* mRNA expression in both bulk tumors and stromal samples (Fig. 5D and E).

**Figure 5.**
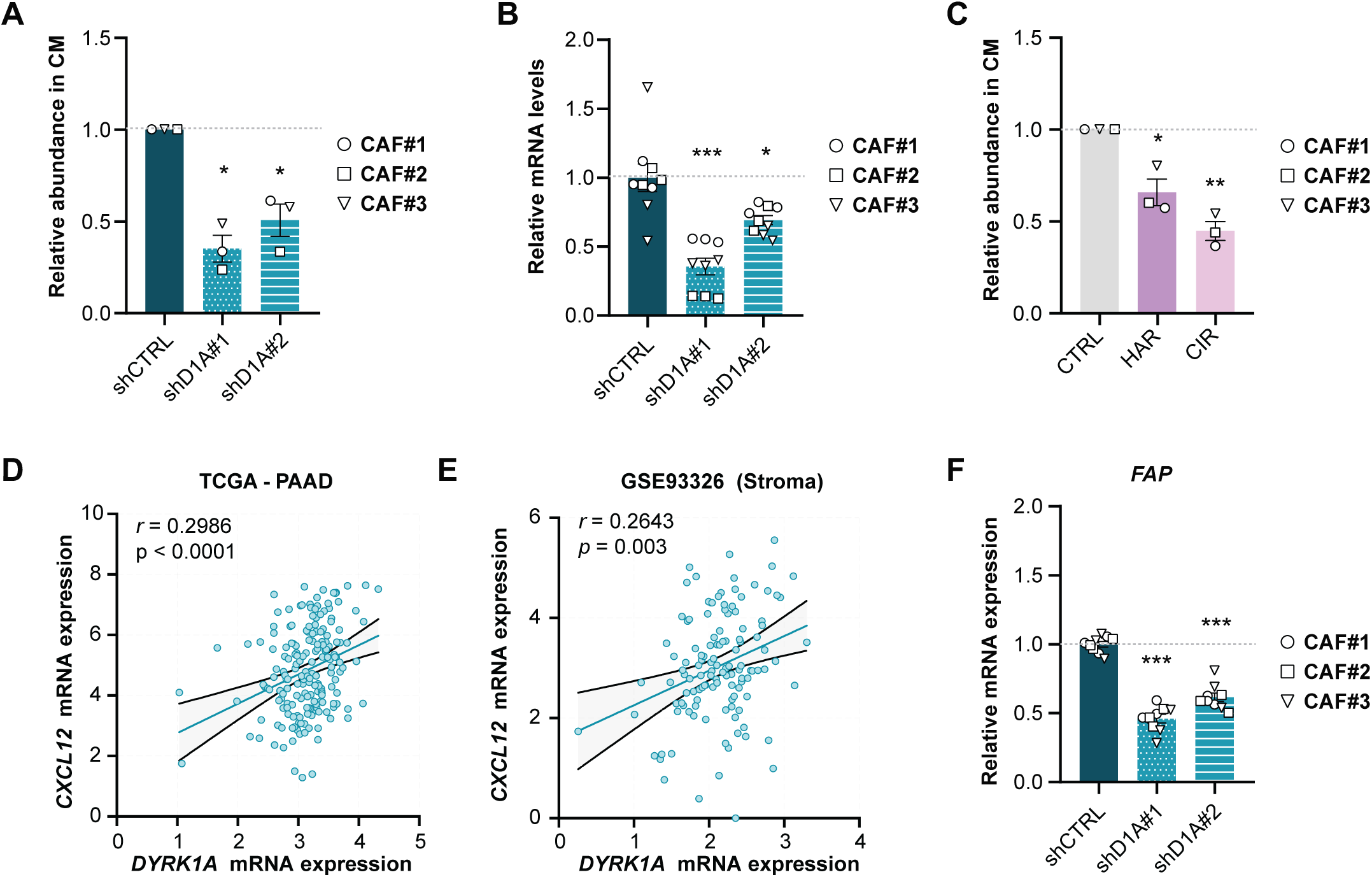
DYRK1A regulates CXCL12 secretion in PDAC CAFs. A, ELISA-based quantification of CXCL12 levels in the CM of shCTRL and shD1A CAF lines. Data are expressed as relative to shCTRL CAFs set as 1 and represented as mean ± SEM. **B,** *CXCL12* mRNA expression analyzed by RT-qPCR in shCTRL and shDYRK1A-expressing CAFs. Data are normalized to the mean expression of shCTRL and represented as mean ± SEM. **C,** ELISA-based quantification of CXCL12 levels in the CM of CAFs treated for 48 h in serum-free conditions with DMSO as vehicle (CTRL), 25 µM harmine (HAR) or 200 nM cirtuvivint (CIR). Data are expressed as relative to vehicle-treated controls set as 1 and represented as mean ± SEM. **D** and **E,** Spearman’s correlation between *DYRK1A* and *CXCL12* mRNA expression in PDAC tumor samples from TCGA (**D**, n = 183) or in the microdissected PDAC stromal compartments (**E**, GSE93326, n = 123). **F,** *FAP* mRNA expression analyzed by RT-qPCR in shCTRL and shD1A CAFs. Data are normalized to the mean expression of shCTRL and represented as mean ± SEM. **A** and **C,** one-sample *t*-test. **B** and **F**, unpaired *t*-test; \**p* ≤ 0.05, \*\**p* ≤ 0.01, \*\*\**p* ≤ 0.001.

In PDAC, CXCL12 is predominantly secreted by fibroblast activation protein (FAP)+ CAFs (24). Consistently, both *CXCL12* and *FAP* expression was markedly higher in the stromal compartment than in epithelial cells (Supplementary Fig. S5C). DYRK1A KD also reduced *FAP* expression in CAFs (Fig. 5F), paralleling the significant reduction detected in two of the three CAF lines by the MS-based secretome analysis (Table S7). Treatment with cirtuvivint lowered *FAP* mRNA levels in all CAF lines (Supplementary Fig. S5D). Supporting these findings, analyses of both bulk TCGA and stromal PDAC datasets revealed a positive correlation between *DYRK1A* and *FAP* expression (Supplementary Fig. S5E and S5F). Overall, these results show that DYRK1A activity modulates the expression of both CXCL12 and FAP in CAFs, pointing toward a coordinated DYRK1A-dependent regulation.

### DYRK1A-dependent chemokine secretion by PDAC cells and CAFs modulates monocyte and MDSC recruitment

The PDAC TME is characterized by abundant immunosuppressive myeloid cells recruited by tumor- and stroma-derived chemokines (4). CCL2, CCL5 and CXCL12 are well-established mediators of monocyte mobilization and MDSCs homing to primary and metastatic PDAC sites, thereby promoting immune evasion (24–27). Additionally, CSF-1 receptor inhibition limits MDSC tumor infiltration (28). Having identified DYRK1A-dependent alterations in the secretion of these factors by PDAC cells and CAFs, we assessed their functional impact using transwell migration assays with human peripheral blood monocytes and *in vitro*-generated monocytic MDSCs. CM from PANC-1 D1A^KO^ clones significantly reduced monocyte chemotaxis compared with WT CM (Fig. 6A), indicating that DYRK1A activity in cancer cells promotes secretion of monocyte-attracting factors. Neutralization of CCL2 in PANC-1 CM diminished monocyte migration (Supplementary Fig. S6A), confirming its contribution. CM from shD1A CAFs also reduced monocyte migration, although to a lesser extent (Fig. 6B).

**Figure 6.**
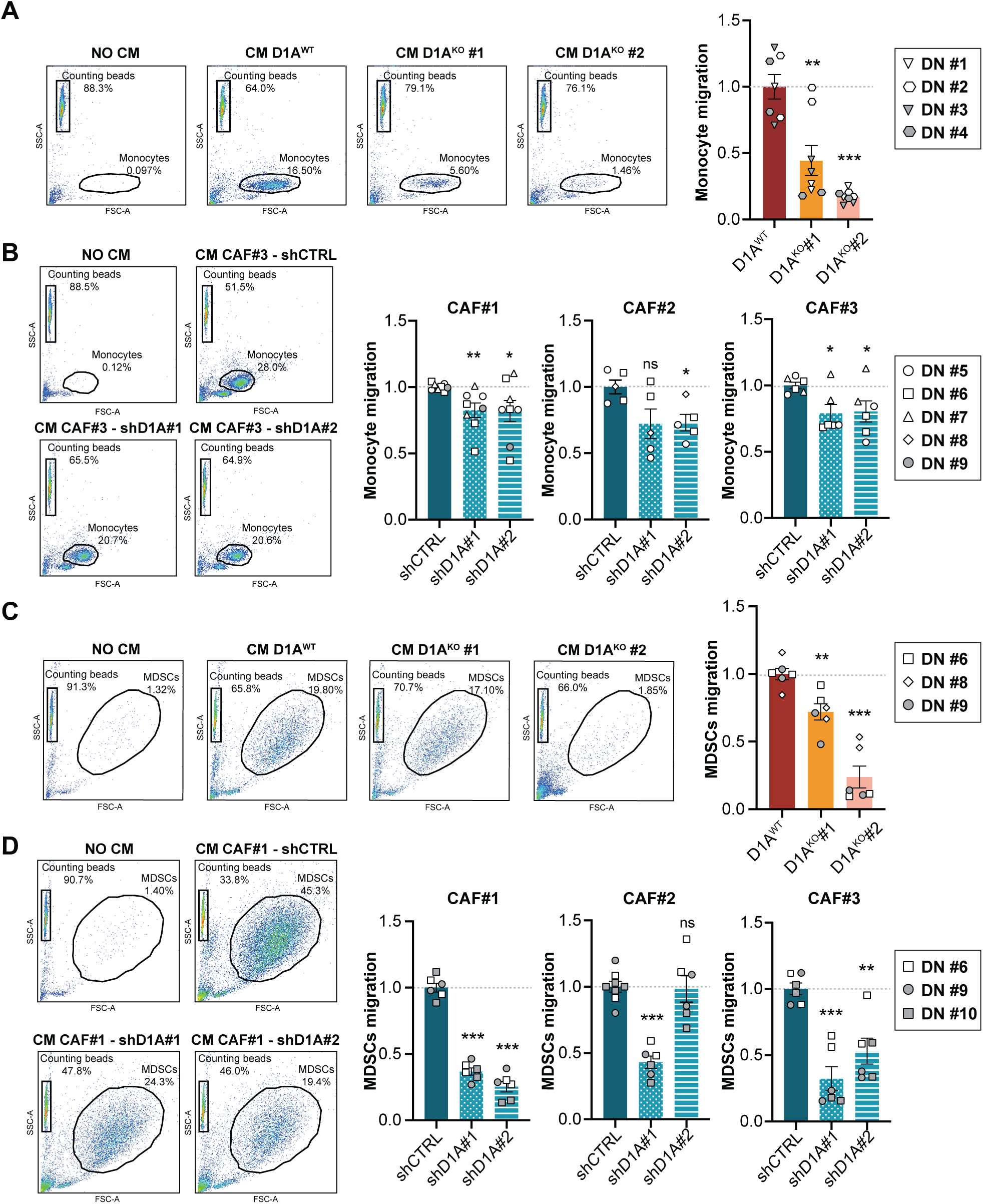
DYRK1A regulates paracrine communication with monocytes and MDSCs. **A** and **B,** Effect of CM from PANC-1 WT or D1A^KO^ clones (**A**), and from shCTRL or shD1A CAFs (**B**) on monocyte migration. Representative flow cytometry plots and quantification from experiments using monocytes from different donors are shown. **C** and **D,** Effect of CM from PANC-1 WT or D1A^KO^ clones (**C**), and from shCTRL or shD1A CAFs (**D**), on the migration of monocytic MDSCs. Representative flow cytometry plots and quantification from experiments using MDSCs from different donors are shown. Migration induced by CM from PANC-1 WT cells (**A** and **C**) or CAFs shCTRL (**B** and **D**) was set as 1. Data are presented as mean ± SEM (*n* = 3-5 donors analyzed with 1-3 replicates; *ns* = not significant, \**p* < 0.05, \*\**p* < 0.01, \*\*\**p* < 0.001; unpaired *t*-test).

We next assessed the impact of the two secretomes on MDSC migration. CM from both PANC-1 D1A^KO^ clones and shD1A CAFs significantly impaired MDSC chemotaxis compared with controls (Fig. 6C and D). Blocking CXCL12 in CAF CM completely abolished MDSC migration, highlighting its role in mediating MDSC attraction by PDAC CAFs (Supplementary Fig. S6B).

Collectively, these results demonstrate that DYRK1A shapes the PDAC secretome to promote the recruitment of immunosuppressive myeloid cells by regulating key chemokines and cytokines in cancer cells and CAFs.

## Discussion

CAFs and cancer cells are major sources of secreted factors that shape the PDAC TME. Dysregulated signals in both compartments can influence tumor progression by altering the secretion of proteins that mediate intercellular communication (4). We previously demonstrated that DYRK1A is overexpressed in PDAC and promotes tumorigenesis by stabilizing c-MET in cancer cells (9). Here, we show that DYRK1A is also highly expressed in CAFs at levels comparable to those in cancer cells. In contrast, DYRK1B expression is minimal in PDAC CAFs, supporting a limited role for this paralogous kinase in the stromal compartment. Whether DYRK1A contributes to CAF activation remains an open question.

Through secretome profiling of DYRK1A-depleted PDAC cancer cells and CAFs, we uncover a previously unrecognized role for this kinase in mediating paracrine interactions within the TME and identify DYRK1A-dependent soluble mediators that affect immune cell infiltration. In PANC-1 cells, DYRK1A depletion broadly altered the secretome, affecting growth factors, cytokines, peptidases and peptidase inhibitors, and ECM components, thus revealing that DYRK1A controls pathways linked to ECM remodeling, cell migration, and PDAC progression. Many DYRK1A-dependent secreted proteins have been proposed as PDAC biomarkers. For instance, MMP7 and CCN2/CTGF, both suggested as diagnostic markers for early disease (19), were downregulated in the secretome of D1A^KO^ PANC-1 clones. As DYRK1A overexpression is detectable in low-grade PanIN lesions (9), early DYRK1A activation might contribute to the secretion of factors associated with disease onset. Beyond diagnostic relevance, several DYRK1A-regulated factors are direct or indirect therapeutic targets, underscoring the potential clinical significance of modulating DYRK1A. In CAFs, proteins upregulated upon DYRK1A KD were enriched in ECM organization. Several of these proteins —including lumican, FSTL1 and SerpinB2/PAI-2— exhibited tumor-restraining roles (29–31), while others such as CHRDL1, IGFBP7, MATN2 and MFAP4 have been associated with improved patient survival (16). Conversely, other upregulated ECM components, like periostin (POSTN) and LOXL2, are linked to PDAC progression (32, 33). Thus, DYRK1A depletion produces a complex ECM secretory profile in CAFs with both pro- and anti-tumorigenic components. This contrasts with PANC-1 D1A^KO^ cells, where ECM pro-tumorigenic proteins were mostly downregulated. Together, these findings point to a complex and context-dependent role of DYRK1A in ECM regulation in PDAC that merits further investigation.

Our study identifies convergent roles for DYRK1A in pancreatic cancer cells and CAFs in regulating the secretion of soluble factors that shape the immune landscape of PDAC TME. DYRK1A depletion or pharmacological inhibition reduced secretion of several immunomodulatory cytokines and chemokines, including CCL2, CCL5, and CSF-1 in cancer cells, and CXCL12 in CAFs. Consistently, public PDAC datasets showed positive correlation between DYRK1A mRNA levels and these mediators, supporting a potential functional link. The distinct transcriptional responses of these DYRK1A-regulated factors suggest that the observed alterations arise from factor-specific mechanisms rather than a single unifying pathway, involving transcriptional and post-transcriptional regulation. Relevant transcription factors include NF-κB/AP-1 and FOXK1 for CCL2 (34, 35), FOXP3 for CCL5 (36), and c/EBPβ and HIF-1 for CXCL12 (37, 38). Post-transcriptional mechanisms such as translational repression of CCL5 (39) have also been described. DYRK1A influences several of these pathways: it promotes non-canonical NF-κB activation (40), represses NF-κB-dependent targets when inhibited (41), interacts with FOXK1 (42), and promotes HIF-1α accumulation (43). Mechanistic studies will thus be needed to elucidate how DYRK1A regulates the secretion of each mediator in epithelial and stromal PDAC cells.

The reduced migration of monocytes and MDSCs toward CM from DYRK1A-depleted PANC-1 cells and CAFs reflects loss of DYRK1A-dependent immunomodulatory signaling. The cytokines and chemokines regulated by DYRK1A —CCL2, CCL5, CSF-1, and CXCL12 — are central mediators of myeloid cell recruitment linked to tumor growth, immunosuppression, and resistance to immune check-point blockade (ICB) in PDAC (4). CCL2 is a paradigmatic myeloid chemoattractant with well-documented effects in PDAC (26, 27). CCL5 and CSF-1 likewise promote monocyte and MDSC infiltration (25, 28). Similarly, CXCL12 drives MDSC accumulation and cytotoxic T-cell exclusion (24, 44), and is enriched in FAP^+^ CAFs, a CAF subtype associated with immune evasion and ICB resistance (24). Consistently, DYRK1A KD or inhibition reduced FAP expression in CAFs alongside CXCL12 downregulation, suggesting a possible role for DYRK1A in maintaining this immunosuppressive CAF population. Further studies will be required to determine whether DYRK1A inhibition preferentially targets FAP^+^ CAFs or modulates CXCL12 secretion independently.

Therapeutic strategies targeting chemokine and cytokine networks are actively explored to overcome immunosuppression and enhance ICB responses in PDAC, including inhibitors of receptors for several DYRK1A-regulated factors (28, 45–47). In this context, our work uncovers a functional link between DYRK1A and chemokine secretion by PDAC cancer cells and CAFs, with implications for the immunosuppressive stroma. Notably, DYRK1A inhibition or deletion increases CD8^+^ T cell infiltration and sensitize tumors to anti-PD-1 therapy in breast cancer and melanoma models (48, 49). DYRK1A may thus emerge as a complementary target to relieve immunosuppression in PDAC by modulating tumor and stroma-derived signals.

In conclusion, with DYRK1A inhibitors now entering clinical development (50), our results provide a rationale for exploring therapeutic strategies that incorporate DYRK1A targeting as part of combinatorial approaches for PDAC treatment.

## Authors’ contributions

**S. Pascual-Sabater:** Investigation, methodology, data curation, manuscript writing. **S. Varhadi:** Investigation, methodology, manuscript writing**. C. Di Vona:** Data curation, formal analysis**. M. Celma:** Investigation, methodology. **S. de la Luna:** conceptualization, data curation, formal analysis, supervision, funding acquisition, manuscript writing and editing**. C. Fillat:** conceptualization, supervision, funding acquisition, manuscript writing and editing.

## Acknowledgements

We thank all the members of Susana de la Luna and Cristina Fillat laboratories for their helpful discussions, and Alicia Raya for technical assistance. We also thank Eva Vaquero for technical help with CAF isolation. We are grateful for the assistance of Eduard Sabidó and Eva Borràs from the CRG/UPF Proteomics Unit of the CRG Core Technologies Programme, which is part of the Spanish Infrastructure for Omics Technologies (ICTS OmicsTech). We are indebted to the Flow Cytometry facility of IDIBAPS for technical help.

