## Supplementary Information for "DYRK1A regulates cancer cell and cancer-associated fibroblast secretomes to foster an immunosuppressive microenvironment in pancreatic cancer"

**Supplementary Data includes**

**Supplementary Materials and Methods**

**Supplementary Table S1:** Chemicals and reagents.

**Supplementary Table S2:** Plasmids.

**Supplementary Table S3:** Oligonucleotides.

**Supplementary Table S4:** Primary and secondary antibodies.

**Supplementary Table S5:** Computational tools and software.

**Supplementary Table S6:** Expression datasets from GEO.

**Supplementary Table S7:** Secreted proteins altered in PANC-1 D1A$^{KO}$ cells and CAFs shD1A, related to Fig. 2 and 3 (Excel file).

**Supplementary Table S8:** Functional enrichment analysis of deregulated secreted proteins in PANC-1 D1A$^{KO}$ cells and CAFs shD1A, related to Fig. 2C, S2F, S2G, 3D, 3E, S3E and S3G (Excel file).

**Supplementary Table S9:** PDAC biomarkers deregulated in PANC-1 D1A$^{KO}$ secretome, related to Fig. 2D (Excel file).

**Supplementary Table S10:** RTKs in the PANC-1 secretome, related to Fig. S2I (Excel file).

**Supplementary Figures**

**Supplementary References**

### Supplementary Materials and Methods

#### Isolation of primary pancreatic fibroblasts

CAFs and normal fibroblasts (NFs) were isolated from human PDAC tissues or adjacent pancreatic tissues, respectively, as described (1) with minor modifications. Briefly, tissues were minced (~1-2 $mm^3$), digested in 250 µg/ml collagenase P and 500 µg/ml DNase I for 15 min at 37°C and further dissociated by pipetting. Suspensions were filtered through a 100-µm nylon mesh, washed with Hanks' Balanced Salt Solution, and fractionated using 12% OptiPrep™. After centrifugation (1,200x*g*, 15 min, no brake), the CAF-enriched pellet was washed and resuspended in 10 ml of DMEM with 10% heat inactivated FBS (iFBS), 15 mM HEPES, 100 units/ml penicillin and 100 µg/ml streptomycin (Pen/Strep), and 2 mM L-glutamine. Fibroblast identity was confirmed by spindle-shaped morphology, *KRAS* exon 2 wild-type status, and expression of fibroblast markers.

#### Lentivirus production and CAF transduction

Lentiviral particles were produced by co-transfecting HEK293FT cells at ~70% confluency with lentiviral constructs along with envelope (pCMV-VSV-G) and packaging (pCMV-ΔR8.91) plasmids using the calcium-phosphate method (CalPhos™ Mammalian Transfection Kit) in the presence of 25 µM chloroquine. Medium was replaced after 16 h, and supernatants harvested at 48 h, filtered (0.45 µm) and stored at -80°C. CAFs were transduced at ~80% confluency in the presence of 8 µg/ml polybrene (hexadimethrine bromide), and fresh medium was added after 24 h.

#### Human monocyte isolation and monocytic MDSCs generation

Peripheral blood monocytes were isolated from buffy coats of healthy donors (Barcelona Public Blood and Tissue Bank) using a Ficoll gradient (Ficoll-Paque Plus density 1.077 g/ml) followed by negative selection. Briefly, 10 ml of buffy coat were diluted to 30 ml with PFE buffer (phosphate-buffered saline (PBS) / 2% FBS / 1 mM EDTA), mixed with 1 ml of RosetteSep™ Human Monocyte Enrichment Cocktail, and incubated for 20 min at room temperature. The mixture was layered onto 15 ml of Ficoll and centrifuged at 1,200x*g* for 25 min (no brake). The monocyte layer was collected, washed twice in PFE buffer (300x*g*, 10 min), and incubated with ACK buffer for erythrocyte lysis. The monocyte-enriched pellet was resuspended in RPMI 1640 medium supplemented with 10% iFBS, 10 mM HEPES, 1% Pen/Strep, and 2 mM L-glutamine. Monocyte purity ($CD14^{+}CD11b^{+}$) exceeded 80% as verified by flow cytometry**.** Monocytes were differentiated into MDSC by culturing with 10 ng/ml IL-6 and 10 ng/ml GM-CSF as described (2).

**Cumulative growth assay**

PANC-1 and derived $DYRK1A^{KO}$ clones were seeded in 24-well plates (1 x $10^4$ cells/well). Cells were counted every other day using an automated cell counter (Countess™, Invitrogen #C10228; RRID:SCR_020149). Growth curves were generated by normalizing cell counts to day 1 values.

**Reverse transcription-quantitative PCR (RT-qPCR)**

Total RNA was isolated using the RNeasy® Plus Kit. Residual genomic DNA was removed with the Turbo DNA-free kit. Reverse transcription was performed with 500 ng RNA and the PrimeScript™ RT Reagent Kit. qPCR was performed in triplicate using specific primers (Supplementary Table S3) and LightCycler® 480 SYBR Green I Master on a QuantStudio™ 7 Pro Real-Time PCR System (Applied Biosystems™, RRID:SCR_020245). Relative mRNA levels were calculated by the $2^{-\Delta\Delta Ct}$ method, normalized to *HPRT1* or *GAPDH*.

**Enzyme-linked immunosorbent assays (ELISA)**

Concentrations of CCL2, CCL5, CSF1 and CXCL12 in CM were determined using DuoSet ELISA kits following manufacturer's instructions. Absorbance at 450 nm ($OD_{450}$) was measured on a BioTek Synergy HT (RRID:SCR_020536) or a Tecan Infinite M200 Pro (RRID:SCR_020543) plate reader. Wavelength correction was performed by subtracting $OD_{540}$, and protein concentrations were calculated from standard curves.

**Western blotting**

Cells were lysed in SDS lysis buffer (25 mM Tris-HCl pH 7.4, 1% SDS, 20 mM β-glycerophosphate, 1 mM EDTA, 10 mM $Na_4P_2O_7$) at 98°C for 20 min. Lysates were centrifuged (16,500x*g*, 5 min, room temperature) and supernatants stored at -20°C. Proteins were resolved by SDS-PAGE and transferred to nitrocellulose or PVDF membranes. The membranes were blocked with 10% non-fat milk for 1 h at room temperature, then incubated with primary antibodies (Supplementary Table S4) overnight at 4°C. Secondary antibodies conjugated to horseradish peroxidase (HRP) or IRDye (Supplementary Table S4) were used for detection. Signals were visualized on a LI-COR-Odyssey imaging system (RRID:SCR_023765) or ImageQuant™ LAS 4000 mini (GE Healthcare, RRID:SCR_018047). Images were processed with Image Studio v5.2.5.

**Immunohistochemistry (IHC)**

Paraffin-embedded 5 µm sections were deparaffinized with xylene and rehydrated through graded ethanol solutions. Antigen retrieval was performed in 10 mM sodium citrate buffer pH 6 for 5 min in a pressure cooker. Sections were blocked for 1.5 h at room temperature with PBS containing 0.3% Triton X-100, 10% FBS, and 1% bovine serum albumin (BSA). After three PBS washes, slides were incubated overnight at 4ºC with primary antibodies (Supplementary Table S4) diluted in PBS-0.1% BSA. Endogenous peroxidase was quenched using Dual Endogenous Enzyme Block (EnVision Detection System) for 10 min. Detection was performed with HRP-conjugated polymer (Agilent #K500711-2) for 30 min, followed by with 3,3-diaminobenzidine (DAB) staining for 5-10 min. Sections were counterstained with

Harris hematoxylin, dehydrated, cleared in xylene, and mounted with DPX Mountant. Imaging was done on an Olympus BX51 microscope (RRID:SCR_018949).

**Sample preparation for mass spectrometry (MS)**

Concentrated CM (10 µg) was reduced with 30 nmol DTT (60 min, 37 ºC) and alkylated in the dark with 60 nmol iodoacetamide. Protein extracts were diluted to 2 M urea in 200 mM ammonium bicarbonate (ABC) and digested sequentially with endoproteinase LysC (1:10 [w:w]; 6 h, 37ºC) and trypsin (1:10 [w/w]; 8 h, 37ºC). Digestion was stopped by addition of 40 µl 100% formic acid (FA). Peptides were desalted using C18 UltraMicroSpin Columns. Briefly, columns were conditioned with 400 µl methanol, equilibrated with 300 µl of 5% FA, and loaded with the digested samples. After two washes with 300 µl of 5% FA, peptides were eluted with 50% acetonitrile/5% FA. Samples were vacuum dried in a SpeedVac Concentrator and reconstituted in 0.1% FA for liquid chromatography coupled to tandem mass-spectrometry (LC-MS/MS) analysis.

**Chromatographic and MS analysis**

Peptides were analyzed using an Orbitrap Fusion Lumos mass spectrometer (Thermo Fisher Scientific, RRID:SCR_020562) coupled to an EASY-nLC 1200 column (Thermo Fisher Scientific, RRID:SCR_014993) at the CRG/UPF Proteomics Unit. Peptides were separated on a 50-cm C18 column (75 µm inner diameter, 2 µm particles, Thermo Fisher Scientific #ES903) using a gradient from 5% to 40% buffer B (0.1% FA in 80% acetonitrile) over 90 min at 300 nl/min.

The mass spectrometer was operated in positive ion mode with nanospray voltage at 2.4 kV and source temperature at 305°C. The acquisition was performed in data-dependent acquisition (DDA) mode and full MS scans with 1 micro scan at resolution of 120,000 were used over a mass range of m/z 350-1400 with detection in the Orbitrap mass analyzer. Auto gain control (AGC) was set to 'standard' and injection time to 'auto'. In each cycle of DDA analysis, following each survey scan, the most intense ions above a threshold ion count of

10,000 were selected for fragmentation. The number of selected precursor ions for fragmentation was determined by the “Top Speed” acquisition algorithm, and a dynamic exclusion of 60 s. Fragment ion spectra were produced via high-energy collision dissociation at normalized collision energy of 28% and acquired in the ion trap mass analyzer in rapid mode. AGC and injection time were set to ‘Standard’ and ‘Dynamic’, respectively, with an isolation window of 1.4 m/z. As quality control, digested BSA standards were run between samples, and instrument performance monitored with QCloud (3).

Spectra were analyzed using Proteome Discoverer v2.5 with Mascot v2.6. Data were searched against the Swiss-Prot human database (April 2022). Parameters for peptide identification were: precursor mass tolerance 7 ppm; fragment ion mass tolerance to 0.5 Da; up to three missed cleavages; carbamidomethylation on cysteine set as fixed modification; oxidation of methionine and N-terminal protein acetylation set as variable modifications. Peptides identified with a FDR < 1% were maintained.

**Computational tools and databases**

RNA expression in pancreatic tumors from The Cancer Genome Atlas Program (TCGA) was obtained at University of California Santa Cruz Xena web site (cohort: GDC TCGA Pancreatic Cancer, Release March 29, 2024) and expressed as $\log_2$ (TPM+1). Single cell RNA-seq analysis of CAFs from different tumor types was performed at a publicly available interactive portal (http://pan-fib.cancer-pku.cn/) (4). A reference secretome list (3,350 proteins) was compiled by integrating UniProt data (release December 2024), selecting entries with “subcellular location [CC]” including the term “secreted” and “Extracellular_Gene Ontology_CC” including “extracellular matrix [GO:0031012]” and “collagen-containing extracellular matrix [GO:0062023]”, together with proteins from the human Surfaceome (5,6) containing a signal peptide predicted by UniProt. Enrichment analysis was performed with EnrichR (August 2025). A list of putative PDAC biomarkers was created with published data (7-9). Therapeutic targets information was obtained from the Therapeutic Structural Antibody

Database (Thera-SAbDab). A full list of computational tools is provided in Supplementary Table S5. The visual representation of data in bar plots and violin plots was done in Prism v8.0.1. Heatmaps, Volcano plots and Sankey plots were generated with the web server SRplot. The comparison and visualization of protein/gene lists in area-proportional Venn diagrams was done with BioVenn. Figures 2A, 3A and S3A were created with BioRender. Microsoft Excel v16.0 was used to create and organize Supplementary Tables S7-S10.

**Supplementary Table S1:** Chemicals and reagents

| Name | Source | Reference |
|---|---|---|
| ACK Lysing Buffer | Gibco™ | A1049201 |
| Amicon™ Ultra-15 10-kDa Centrifugal Filter Units | Millipore | 10781543 |
| Ammonium bicarbonate (ABC) | Sigma-Aldrich | 09830 |
| BSA, Trypsin-digested MS Standard (CAM-modified) | New England Biolabs | P8108S |
| CalPhos™ Mammalian Transfection Kit | TakaraBio | 631312 |
| Cirtuvivint | MedChemExpress | HY-137435 |
| Citrate Buffer pH 6 | Sigma-Aldrich | C9999 |
| Collagenase P | Roche | 11213865001 |
| CountBright™ Absolute Counting Beads | Invitrogen | C36950 |
| DMEM Cell Culture Media | Gibco™ | 419660 |
| DMSO | Sigma-Aldrich | D2650 |
| DNase I | Roche | 11284932001 |
| DPX Mountant | Sigma-Aldrich | 06522 |
| ELISA DuoSet Human CCL2/MCP-1 kit | R&D Systems | DY279 |
| ELISA DuoSet Human CCL5/RANTES kit | R&D Systems | DY278 |
| ELISA DuoSet Human CSF1/M-CSF kit | R&D Systems | DY216 |
| ELISA DuoSet Human CXCL12/SDF-1 kit | R&D Systems | DY350 |
| EnVision Detection System, Peroxidase/DAB, Rabbit/Mouse, HRP | Agilent | K500711-2 |
| Fetal Bovine Serum | Gibco™ | 10270106 |
| Ficoll-Paque Plus Density Gradient Media | Cytiva | 17144003 |
| L-glutamine | Gibco™ | 25030081 |
| GM-CSF, Human Recombinant | R&D Systems | 215-GM |
| Harmine | Sigma-Aldrich | 286044 |
| Hanks' Balanced Salt Solution | Gibco™ | 14025092 |
| Hematoxylin | PanReac AppliChem | 256991 |
| Interleukin 6, Human Recombinant | Peprotech | 200-06 |
| Iodoacetamide | Sigma-Aldrich | I1149 |
| LightCycler® 480 SYBR Green I Master | Roche | 04707516001 |
| Lipofectamine 3000 | Invitrogen | L3000015 |
| Lys-C endopeptidase | Wako | 129-02541 |
| Nonfat Dry Milk | Cell Signaling Technologies | 9999S |
| OptiPrep™ Density Gradient Media | Sigma-Aldrich | D1556 |
| Penicillin-Streptomycin Solution | Gibco™ | 15140130 |
| Polybrene | Sigma-Aldrich | H9268 |
| PrimeScript™ RT Reagent Kit | Takara Bio | RR037A |
| Puromycin | Sigma-Aldrich | P8833 |
| RosetteSep™ Human Monocyte Enrichment Cocktail | STEMCELL Technologies | 15028 |
| RNeasy® Plus Mini Kit | Qiagen | 74134 |
| RPMI 1640 Cell Culture Media | Gibco™ | 61870010 |
| Trypsin, Sequencing Grade Modified | Promega | V5113 |
| Turbo DNA-free kit | Invitrogen | AM1907 |
| UltraMicroSpin Columns, C18 stage | The Nest Group INC | SS18V |

| | | |
|---|---|---|
| Urea | Cytiva | 17-1319-01 |
| Western blotting Membranes, Nitrocellulose Amersham™ Protran® Premium | Sigma-Aldrich | 10600003 |
| Western blotting Membranes, Immobilon®-P, PVDF | Millipore | IPVH20200 |

**Supplementary Table S2:** Plasmids

| Reagent | Source | Identifier |
|---|---|---|
| 2X_pX458_pSpCas9(BB)-2A-GFP | Addgene (10) | RRID:Addgene_172221 |
| pCMV-ΔR8.91 | (11) | N/A |
| pCMV-VSV-G | Addgene (12) | RRID:Addgene_8454 |
| pLKO.1-puro shRNA Control Plasmid DNA (shCTRL) | Sigma-Aldrich | SHC002 |
| pLKO.1-puro-shD1A#1 | Sigma-Aldrich | TRCN0000022999 |
| pLKO.1-puro-shD1A#2 | Sigma-Aldrich | TRCN0000199188 |
| pLOX-Ttag-iresTK | Addgene (13) | RRID:Addgene_12246 |

**Supplementary Table S3:** Oligonucleotides

| Gene | | Primer sequence (5'->3') | Application |
|---|---|---|---|
| ***CCL2*** | Forward | ATCACCAGCAGCAAGTGTC | RT-qPCR |
| | Reverse | AGGTGGTCCATGGAATCCTG | |
| ***CCL5*** | Forward | GAAGGAAGTCAGCATGCCTCTA | RT-qPCR |
| | Reverse | CATGTTTGCCAGTAAGCTCCTG | |
| ***CXCL12*** | Forward | CTCAACACTCCAAACTGTGCCC | RT-qPCR |
| | Reverse | CTCCAGGTACTCCTGAATCCAC | |
| ***DYRK1A*** | Forward | GCTGGACATCCAACATACCA | RT-qPCR |
| | Reverse | TCTGTTGCACACAAACTCCTG | |
| ***DYRK1A*** | D1A$^{WT}$_fwd | GGCATATGATCGTGTGGAGC | DYRK1A$^{KO}$ genotyping |
| | D1A$^{KO}$_fwd | TCTGCTAGTTGTGAACCTGGT | |
| | D1A_rev | TTCCTTATGCTTTTCTTCCATCA | |
| ***DYRK1B*** | Forward | CGAGCAGTTTGAGTCCCCTT | RT-qPCR |
| | Reverse | ATCAGGCAATACCTGCGTGT | |
| ***DYRK1B*** | D1B$^{WT}$_fwd | CTGTCCTCCTTTCCCTGTGA | DYRK1B$^{KO}$ |
| | D1B$^{WT}$_rev | AGAATGTCTTGGGCAACCAC | genotyping |
| | D1B$^{KO}$_fwd | CACCCCTTTCTTCGTGACAT | DYRK1B$^{KO}$ |
| | D1B$^{KO}$_rev | CACCCCTTTCTTCGTGACAT | genotyping |
| ***FAP*** | Forward | GGAAGTGCCTGTTCCAGCAATG | RT-qPCR |
| | Reverse | TGTCTGCCAGTCTTCCCTGAAG | |
| ***GAPDH*** | Forward | TGTCAAGCTCATTTCCTGGTATGA | RT-qPCR |
| | Reverse | TTACTCCTTGGAGGCCATGTGGG | |
| ***HPRT1*** | Forward | GATATAAGCCAGACTTTGTTGGATTTG | RT-qPCR |
| | Reverse | CTTGAACTCTCATCTTAGGCTTTG | |
| ***KRAS*** | Forward | GGTGGAGTATTTGATAGTGTA | CAF genotyping |
| | Reverse | GGTCCTGCACCAGTAATATGC | |

**Supplementary Table S4:** Primary antibodies

| **Antibodies** | **Vendor (catalog #); RRID** | **Assay (dilution)** |
|---|---|---|
| ß-Actin | Sigma-Aldrich (#A2066); RRID:AB_476693 | WB (1:10,000) |
| CCL2 | R&D Biotechne (#MAB279); RRID:AB_2071645 | Neutralization (4 µg/mL) |
| CD11b-APC | Biolegend (#101212); RRID:AB_312795 | Flow cytometry (1:50) |
| CD14 | BD Biosciences (#566141); RRID:AB_2739539 | Flow cytometry (1:100) |
| CK19 | Abcam (#ab52625); RRID:AB_2281020 | IHC (1:500) |
| CXCL12 | R&D Biotechne (#MAB310); RRID:AB_2276927 | Neutralization (100 µg/mL) |
| DYRK1A | Santa Cruz (#sc-100376); RRID:AB_1122375 | WB (1:500) |
| DYRK1A | Abnova (#H00001859-M01); RRID:AB_534844 | IHC (1:200) |
| DYRK1B | Abcam (#ab113968); RRID:AB_10865501 | WB (1:500) |
| GAPDH | Sigma-Aldrich (#ABS16); RRID:AB_10806772 | WB (1:2,500) |
| □-Tubulin | Sigma-Aldrich (#T6199); RRID:AB_477583 | WB (1:10,000) |
| Vimentin | Leica (#NCL-L-VIM-V9); RRID:AB_564055 | IHC (1:50) |

**Secondary antibodies for WB**

| **Secondary antibodies** | **Vendor (catalog #); RRID** | **Dilution** |
|---|---|---|
| Donkey anti-mouse IRDye 800CW | LICOR (#926-32212); RRID:AB_621847 | 1:10,000 |
| Donkey anti-rabbit IRDye 680RD | LICOR (#926-68073); RRID:AB_10954442 | 1:10,000 |
| Rabbit anti-mouse HRP-conjugated | Agilent (#P0260); RRID:AB_2636929 | 1:2,000 |
| Goat anti-rabbit HRP-conjugated | Agilent (#P0448); RRID:AB_2617138 | 1:2,000 |

**Supplementary Table S5:** Computational tools and software

| Name | Source | Identifier |
|---|---|---|
| BioVenn | https://biovenn.nl (14) | RRID:SCR_026853 |
| BioRender | http://biorender.com | RRID:SCR_018361 |
| EnrichR | https://maayanlab.cloud/Enrichr (15) | RRID:SCR_001575 |
| Excel v16.0 | Microsoft | RRID:SCR_016137 |
| FlowJo™ v10 | Becton Dickinson | RRID:SCR_008520 |
| GEO database | https://www.ncbi.nlm.nih.gov/geo (16) | RRID:SCR_005012 |
| Image Studio v5.2.5 | LI-COR | RRID:SCR_015795 |
| Mascot Search Engine v2.6 | Matrix Science | RRID:SCR_014322 |
| Prism v8.0.1 | GraphPad Software | RRID:SCR_002798 |
| Proteome Discoverer v2.5 | Thermo Fisher Scientific | RRID:SCR_014477 |
| SC Fibroblast Atlas | http://pan-fib.cancer-pku.cn (4) | N/A |
| SRplot | https://www.bioinformatics.com.cn/en | RRID:SCR_025904 |
| Thera-SAbDab | http://opig.stats.ox.ac.uk/webapps/therasabdab (17) | RRID:SCR_022093 |
| UniProt | https://www.uniprot.org (18) | RRID:SCR_002380 |

**Supplementary Table S6:** Expression datasets from GEO

| Accession | Information | Reference |
|---|---|---|
| GSE93326 | high throughput sequencing of epithelium and stroma RNA samples from human PDAC frozen sections | (19, 20) |
| GSE164665 | laser capture microdissection of PDAC samples | (21) |
| GSE71729 | expression profiling by array of gene expression in PDAC | (22) |

### Supplementary Figures

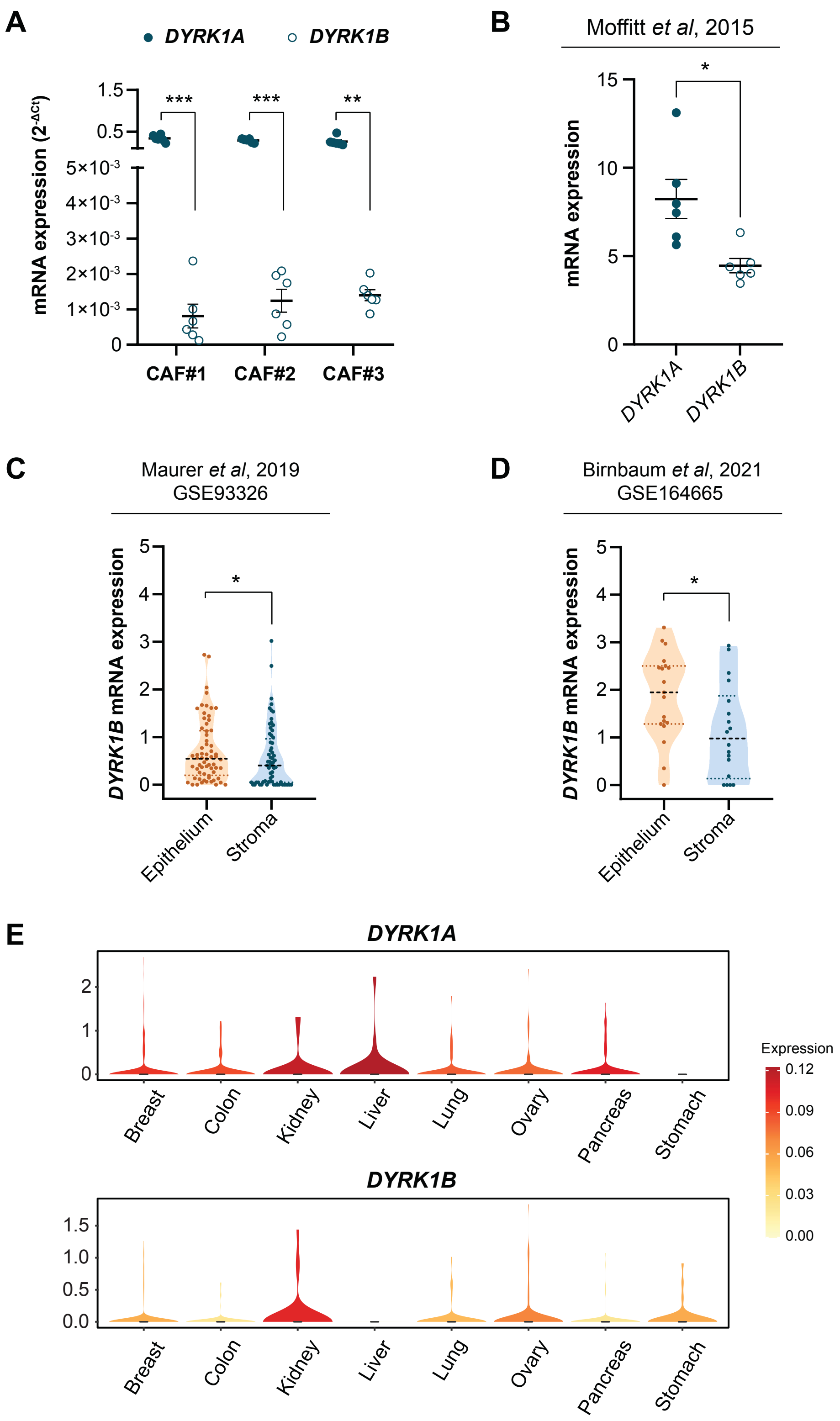


**Figure S1. Accompanying data for Figure 1. A,** *DYRK1A* and *DYRK1B* mRNA expression analyzed by RT-qPCR in CAFs from 3 PDAC patients. *HPRT1* was used as a housekeeping gene. Data are represented as mean ± SEM (*n* = 6). **B,** *DYRK1A* and *DYRK1B* mRNA expression (FPKM) in CAFs from 6 different PDAC patients as reported in Moffit et al., 2015 (22). Data are represented as mean ± SEM (*n* = 6). **C** and **D,** Violin plots showing differential *DYRK1B* expression ($\log_2$ [TPM+1]) in matched epithelial and stromal compartments of microdissected PDAC tumors from GSE93326 (**C,** *n* = 64) or GSE164665 (**D**, *n* = 18). **E,** scRNA-seq data of *DYRK1A* and *DYRK1B* expression in fibroblasts from the indicated tumor types. Violin plots were generated using the webtool developed by Gao et al., 2024 (4) (http://pan-fib.cancer-pku.cn/).

**A** and **B**, two-tailed paired *t*-test; **C** and **D**; Wilcoxon matched-pairs signed rank test; *ns* = not significant, **p* ≤ 0.05, ***p* ≤ 0.01; ****p* ≤ 0.001.

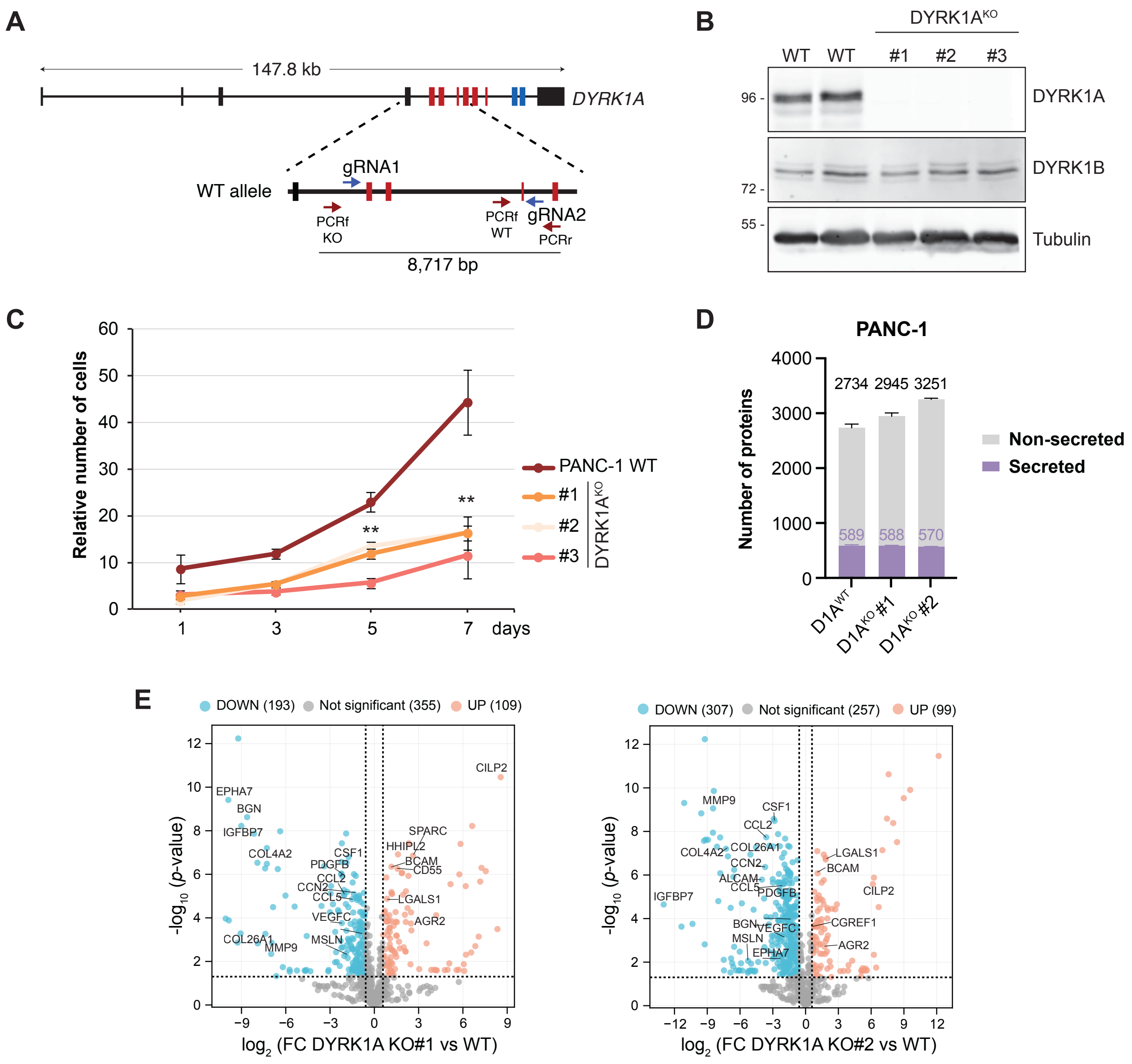
A
147.8 kb
DYRK1A
WT allele
gRNA1
PCRf KO
PCRf WT
gRNA2
PCRr
8,717 bp
B
DYRK1A^KO
WT WT #1 #2 #3
96 -
72 -
55 -
DYRK1A
DYRK1B
Tubulin
C
Relative number of cells
days
PANC-1 WT
#1
#2
#3
DYRK1A^KO
**
**
D
PANC-1
Number of proteins
2734 2945 3251
589 588 570
Non-secreted
Secreted
D1A^WT
D1A^KO #1
D1A^KO #2
E
DOWN (193) Not significant (355) UP (109)
-log10 (p-value)
log2 (FC DYRK1A KO#1 vs WT)
DOWN (307) Not significant (257) UP (99)
log2 (FC DYRK1A KO#2 vs WT)
EPHA7
BGN
IGFBP7
COL4A2
CSF1
PDGFB
CCL2
CCN2
CCL5
VEGFC
COL26A1
MMP9
MSLN
CILP2
SPARC
HHIPL2
BCAM
CD55
LGALS1
AGR2
ALCAM
CGREF1

F

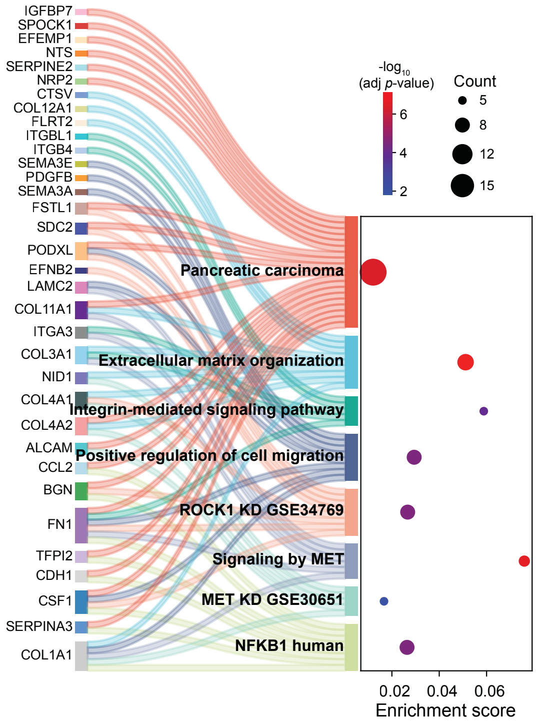


G

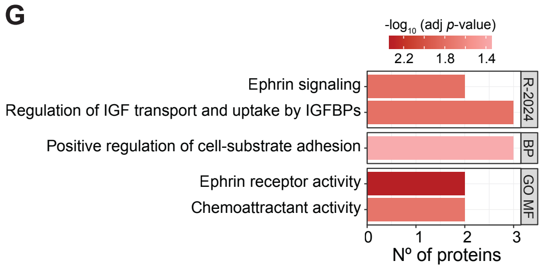


H

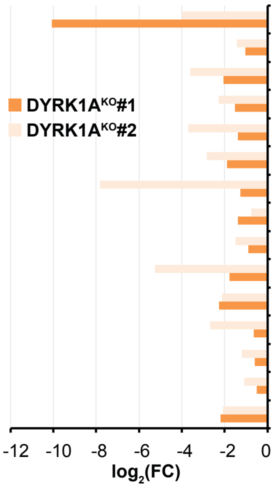


| Secretome | Therapeutic | Disease | Receptor/ Ligand | Therapeutic | Disease |
|---|---|---|---|---|---|
| **ALCAM** | Praluzatamab | Solid tumors | - | - | - |
| **AXL** | Mecbotamab | Solid tumors; Soft tissue sarcoma | - | - | - |
| **CCL2** | Carlumab | Solid tumors | **CCR2** | Plozalizumab | Solid tumors |
| **CCL5** | - | - | **CCR5** | Leronlimab | Solid tumors |
| **CCN2** | Pamrevlumab | Pancreatic cancer | - | - | Autoimmune disorders |
| **CSF1** | Lacnotuzumab | Solid tumors | **CSF1R** | Emactuzumab | Pancreatic cancer |
| **EBI3 (IL27)** | Casdozokitug | Solid tumors | - | - | - |
| **IGF2** | Xentuzumab | Breast cancer; Prostate cancer | - | - | - |
| **LOXL2** | Simtuzumab | Pancreatic cancer | - | - | - |
| **MSLN** | Misitatug | Solid tumors | - | - | - |
| **PDGFB** | - | - | **PDGFRA** | Olaratumab | Pancreatic cancer |
| **ROR1** | Zilovertamab | CLL; Prostate cancer; Endometrial cancer; Ovarian cancer | | | |
| **TGFB2** | Nisevokitug | Pancreatic cancers; Colorectal cancers; Myelodysplastic syndromes | | | |
| **VEGFA** | Suvemcitug | Solid tumors | **VEGFR2** | Olinvacimab | Solid tumors |
| **VEGFC** | - | - | **VEGFR2** | Pulocimab | Solid tumors |

I

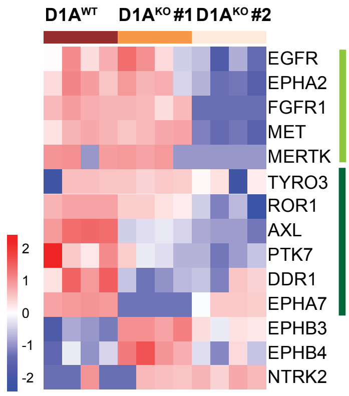

**Figure S2. Accompanying data for Figure 2. A,** CRISPR-Cas9 approach for DYRK1A knockout in PDAC cell lines. Schematic representation of the genomic structure of human *DYRK1A*: boxes represent exons (in red, those encoding the catalytic domain) and lines represent introns. The gene size is indicated. The region deleted by CRISPR/Cas9 is magnified, with blue arrows indicating the position of gRNAs and red arrows indicating the primers used to confirm the deletion. **B,** Western blot analysis of DYRK1A and DYRK1B expression in PANC-1 $D1A^{KO}$ clones. Tubulin was used as a loading control. **C,** Cumulative growth curves of PANC-1 WT cells and $D1A^{KO}$ clones. Equal number of cells were seeded at day 0 and counted at the indicated time points. The graphs show a representative experiment (mean ± SEM of 3 independent replicates) of two performed. Results are expressed relative to the number of seeded cells. Differences in cell numbers were statistically significant at days 5 and 7 for all clones (**$p \leq 0.01$; unpaired *t*-test). **D**, Number of proteins detected in the CM of PANC-1 WT cells and derived $D1A^{KO}$ clones represented as mean ± SEM of the four replicates analyzed by MS. Bars are subdivided to indicate proteins classified as secreted (purple) or non-secreted (gray) according to our annotation pipeline. The number of proteins (total and secreted) detected in each condition is shown above the stacked bars. **E,** Volcano plots showing differentially secreted proteins between PANC-1 WT and $D1A^{KO}$ clones, with significance ($-\log_{10}$ *p*-value) plotted against differences in protein abundance $\log_2(FC)$. The dotted line along the *y*-axis indicates significance ($p \leq 0.05$), while vertical dotted lines indicate fold change thresholds ($\log_2(FC) \geq 0.5$ or $\leq -0.5$). **F,** Sankey plot showing the association between downregulated secreted proteins common to both PANC-1 $D1A^{KO}$ clones and their enriched functional categories, with links representing protein-to-pathway assignments. **G,** Functional enrichment analysis of secreted proteins commonly upregulated in both PANC-1 $D1A^{KO}$ clones (27 proteins). All significantly enriched terms in the categories indicated are shown. R-2024, Reactome Pathways 2024; BP, GO Biological Processes 2025, GO MF, GO Molecular Functions 2025. Extended data is available in Table S8. **H**, Extended information for Fig. 2E. Bar graphs show $\log_2(FC)$ in protein abundance between PANC-1 $D1A^{KO}$ clones and WT cells, along with information on available therapeutic antibodies and their disease indications (ligand- or receptor-targeting, as indicated). **I**, Heatmap showing RTKs detected in the CM of PANC-1 WT and D1A$^{KO}$ clones. The light green bar indicates RTKs with reduced expression in one D1A$^{KO}$ clone, while the dark green bar highlights RTKs with reduced expression in both clones (extended information in Table S10).

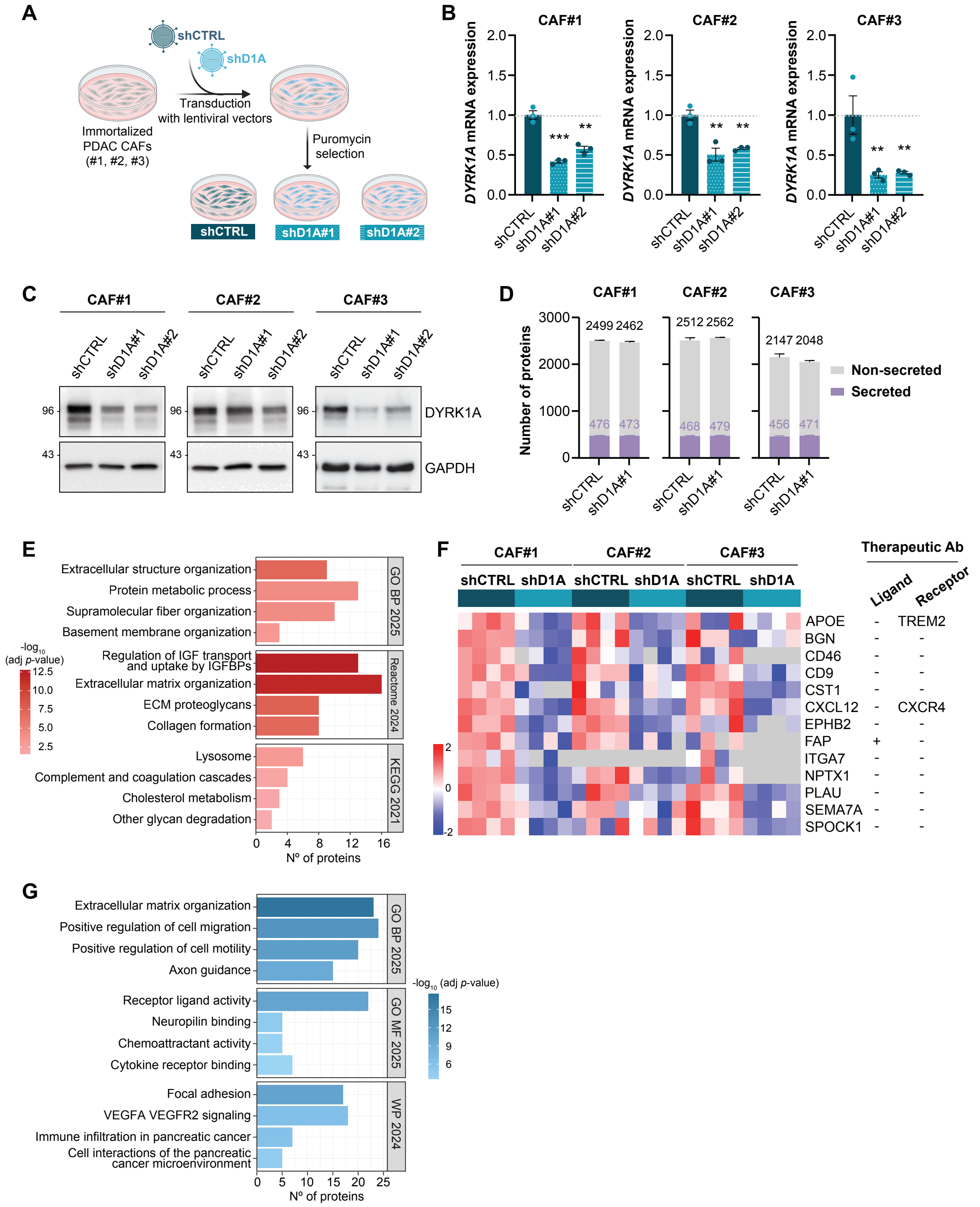

**Figure S3. Accompanying data for Figure 3. A,** Experimental scheme of the knockdown strategy used to generate PDAC CAFs with reduced DYRK1A expression. **B,** Analysis of *DYRK1A* expression by RT-qPCR in shCTRL and shD1A PDAC CAFs. *HPRT1* was used as a housekeeping gene. Data are normalized to the mean expression of shCTRL and represented as mean ± SEM ($n$ = 3; *$p \leq 0.05$, **$p \leq 0.01$, ***$p \leq 0.001$; unpaired $t$-test). **C,**

Western blot analysis of DYRK1A expression in shCTRL and shD1A PDAC CAFs. GAPDH was used as a loading control. **D,** Number of proteins identified in the CM of shCTRL and shD1A CAFs, represented as mean ± SEM of the four replicates analyzed by MS. Bars are subdivided to indicate proteins classified as secreted (purple) or non-secreted (gray) according to our annotation pipeline. The number of proteins (total and secreted) detected in each condition is shown above the stacked bars. **E,** Functional enrichment analysis of upregulated proteins detected in the CM of shD1A CAF lines (shared by at least two CAF lines, 73 proteins). Selected terms from Gene Ontology Biological Process 2025 (GO BP 2025), Reactome Pathways 2024 (Reactome 2024), and Kyoto Encyclopedia of Genes and Genomes 2021 (KEGG 2021) are shown. Extended data is available in Table S8. **F**, Heatmap ($log_2$-normalized abundance, *z*-scored by row and CAF line) showing downregulated proteins in the CM of shD1A CAFs. The association with existing therapeutic antibodies targeting either the ligand or the receptor is indicated. TREM2 receptor for APOE: Iluzanebart, active for Leukoencephalopathies and Neurodegenerative disorders; CXCR4, receptor for CXCL12: Ulocuplumab in clinical trials for acute myeloid leukemia, solid tumors, Waldenstrom's macroglobulinemia and neuroectodermal tumors; FAP: Simlukafusp for malignant melanoma, renal cell carcinoma, solid tumors (discontinued). **G**, Functional enrichment analysis of downregulated proteins common in DYRK1A-depleted PANC-1 cells and CAF lines. Proteins shared by the two PANC-1 D1A$^{KO}$ clones and detected in at least two shD1A CAF lines were included in the analysis (157 proteins). Selected terms from GO BP 2025, Gene Ontology Molecular Function 2025 (GO MF 2025), and WikiPathways 2024 Human (WP 2024) are shown. Extended data is available in Table S8.

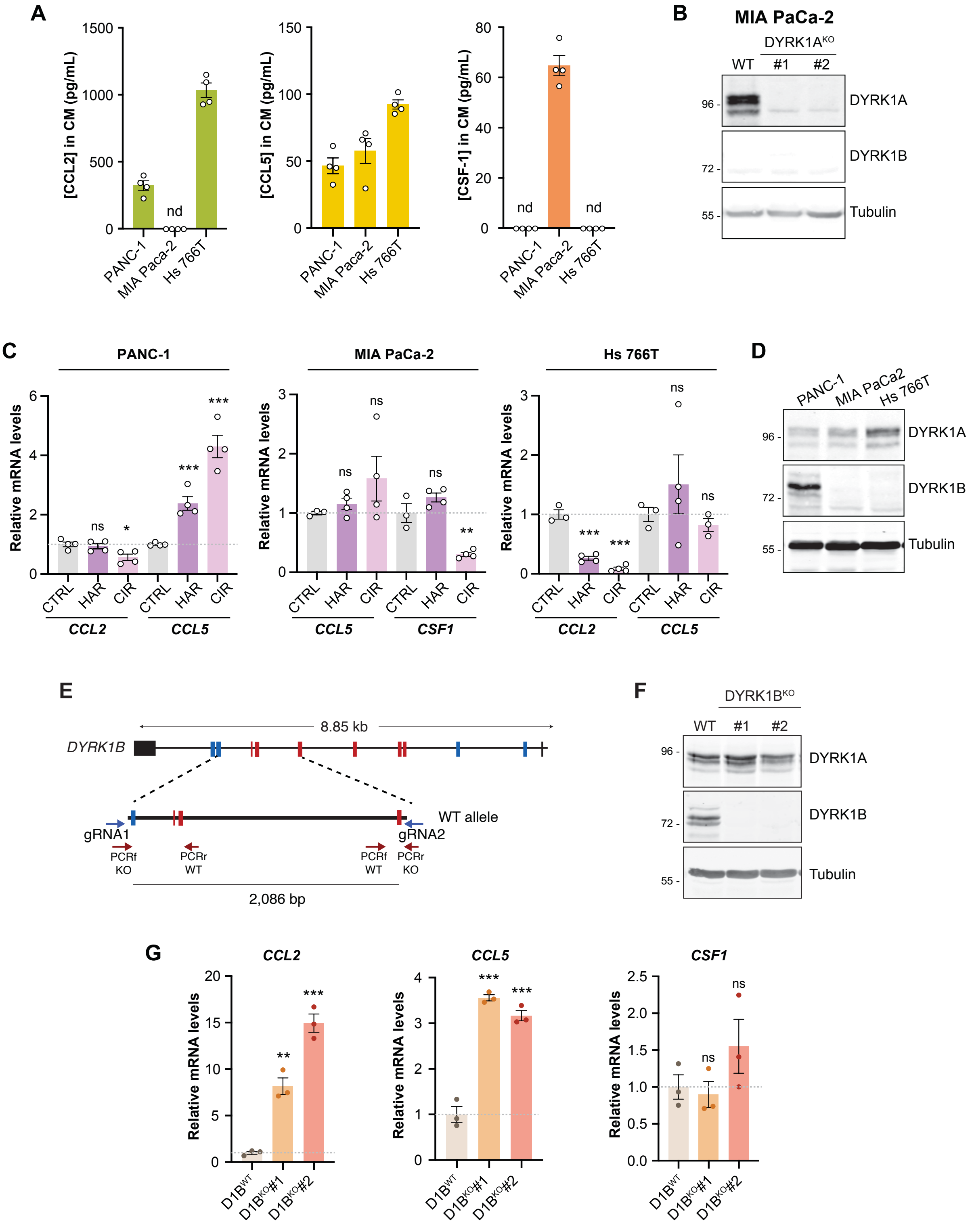
A
[CCL2] in CM (pg/mL)
[CCL5] in CM (pg/mL)
[CSF-1] in CM (pg/mL)
nd
PANC-1
MIA Paca-2
Hs 766T
B
MIA PaCa-2
DYRK1AKO
WT
#1
#2
DYRK1A
DYRK1B
Tubulin
C
PANC-1
MIA PaCa-2
Hs 766T
Relative mRNA levels
CTRL
HAR
CIR
CCL2
CCL5
CSF1
ns
D
PANC-1
MIA PaCa2
Hs 766T
E
DYRK1B
8.85 kb
WT allele
gRNA1
gRNA2
PCRf KO
PCRr WT
PCRf WT
PCRr KO
2,086 bp
F
DYRK1BKO
G
D1BWT
D1BKO#1
D1BKO#2

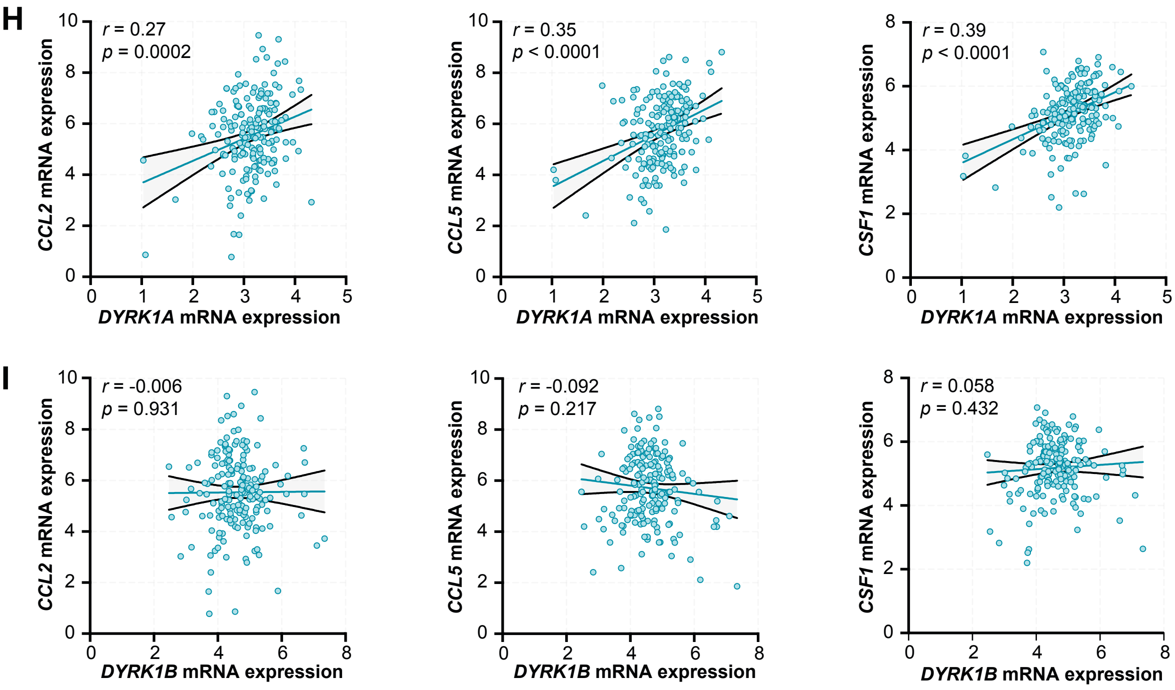


**Figure S4. Accompanying data for Figure 4. A,** ELISA-based analysis of CCL2, CCL5 and CSF-1 levels (in pg/mL) in the CM of PDAC cell lines. **B,** Western blot analysis showing expression of DYRK1 kinases in MIA PaCa-2 D1A$^{KO}$ clones. Tubulin was used as loading control. **C,** mRNA levels of *CCL2*, *CCL5* and *CSF1* in PANC-1, MIA PaCa-2 and Hs 766T cells treated for 24 h in serum-free conditions with DMSO as vehicle (CTRL), 10 μM harmine (HAR) or 1 μM cirtuvivint (CIR). Values are expressed relative to vehicle-treated controls and represented as mean ± SEM. **D,** Western blot showing expression of DYRK1 kinases in the PDAC cell lines indicated. Tubulin was used as loading control. **E**, CRISPR-Cas9 approach for DYRK1B inactivation in PANC-1 cells. Schematic representation of the genomic structure of human *DYRK1B*: boxes represent exons (in red, those encoding for the catalytic domain) and lines represent introns. The gene size is indicated. The region deleted by CRISPR/Cas9 is magnified, with blue arrows indicating the position of the gRNA target sites and red arrows indicating the primers used to confirm the deletion. **F,** Western blot showing expression of DYRK1 kinases in PANC-1 D1B$^{KO}$ clones. Tubulin was used as loading control. **G,** mRNA expression of *CCL2*, *CCL5* and *CSF1* analyzed by RT-qPCR in PANC-1 cells and D1B$^{KO}$

clones. *HPRT1* was used as a housekeeping gene. Data are expressed relative to parental PANC-1 cells (WT) values and represented as mean ± SEM. **H** and **I,** Spearman's correlation analysis showing gene expression levels ($log_2$ [TPM+1]) of *DYRK1A* (**I**) or *DYRK1B* (**J**) versus *CCL2*, *CCL5* or *CSF1* in TCGA PDAC samples (n =183). Spearman's correlation coefficient (*r*) and *p*-values are shown for each analysis.

In panels **C** and **G**: *nd* = not detected, *ns* = not significant, $*p \leq 0.05$, $**p \leq 0.01$, $***p \leq 0.001$; unpaired *t*-test.

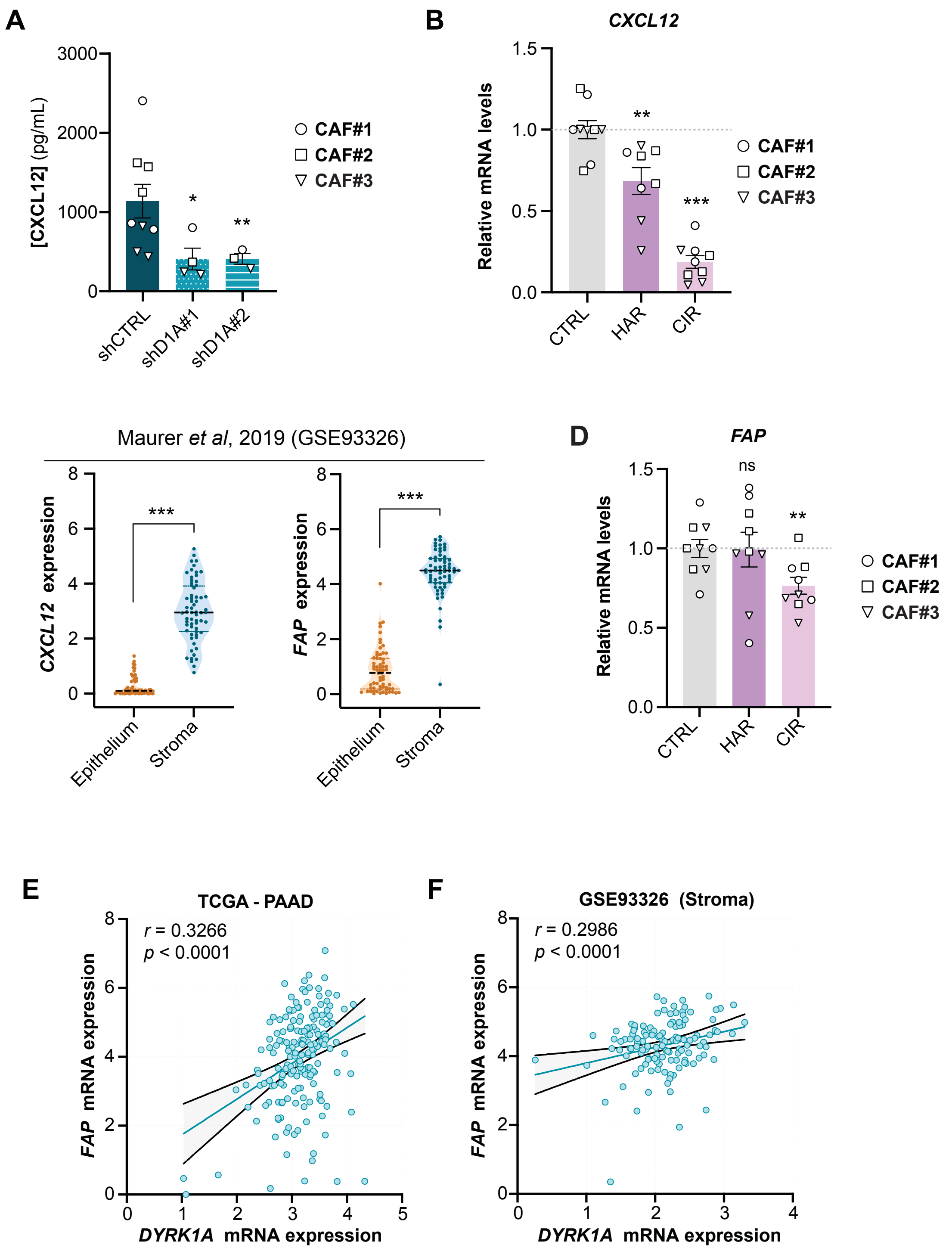


**Figure S5. Accompanying data for Figure 5. A,** CXCL12 concentration (pg/mL) in the CM of shCTRL and shD1A measured by ELISA. **B** and **D,** mRNA expression of *CXCL12* (**B**) and *FAP* (**D**) analyzed by RT-qPCR in CAFs treated for 24 h in serum-free conditions with DMSO as vehicle (CTRL), 25 µM harmine (HAR) or 200 nM cirtuvivint (CIR). Data are normalized by

*HPRT1* expression in individual experiments and represented as mean ± SEM ($n$ = 2-3 per CAF line). **C**, Violin plots showing *CXCL12* and *FAP* mRNA expression in matched epithelial and stromal compartments of microdissected PDAC tumors (GSE93326, $n$ = 64). **E** and **F,** Spearman's correlation analysis showing gene expression levels ($log_2$ [TPM+1]) of *DYRK1A* and *FAP* in TCGA PDAC samples (**E**, n = 183) or in the microdissected PDAC stromal compartment (**F**, GSE93326, n = 123). Spearman's correlation coefficient ($r$) and $p$-values are shown for each analysis.

**A**, unpaired $t$-test with Welch's correction; **B** and **D**; unpaired $t$-test; **C**, Wilcoxon matched-pairs signed rank test; *ns* = not significant, *$p \leq 0.05$, **$p \leq 0.01$; ***$p \leq 0.001$.

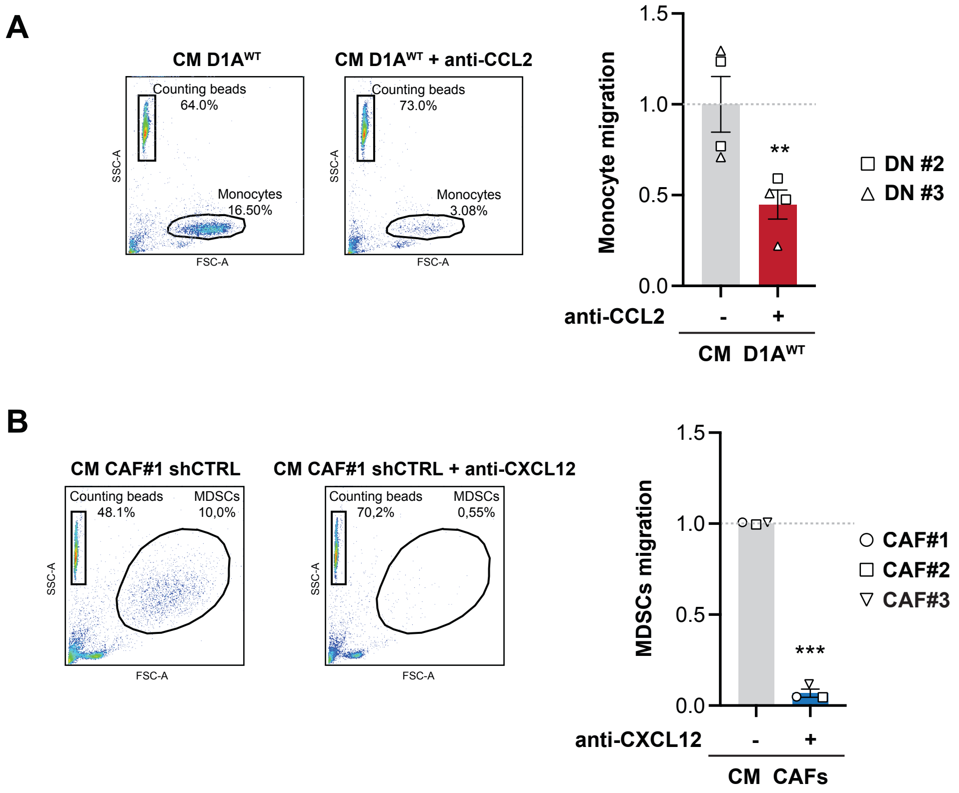


**Figure S6. Accompanying data for Figure 6. A,** Effect of CCL2 neutralization on monocyte migration induced by CM from PANC-1 WT cells. Representative plots and quantification of the results ($n$ = 2 donors analyzed in duplicate; $^{**}p \leq 0.01$; unpaired $t$-test). **B**, Effect of CXCL12 neutralization on MDSCs migration induced by CM from CAFs. Representative plots and quantification of the results using CM from different CAFs on MDSCs from one donor are shown ($^{***}p < 0.001$; unpaired $t$-test).
